# Cancer associated fibroblasts drive epithelial to mesenchymal transition and classical to basal change in pancreatic ductal adenocarcinoma cells with loss of IL-8 expression

**DOI:** 10.1101/2025.01.07.631784

**Authors:** Samantha Guinn, Brayan Perez, Joseph A. Tandurella, Mili Ramani, Jae W. Lee, Daniel J. Zabransky, Emma Kartalia, Jignasha Patel, Haley Zlomke, Norman Nicolson, Sarah Shin, Benjamin Barrett, Nicholas Sun, Alexei Hernandez, Erin Coyne, Courtney Cannon, Nicole E. Gross, Soren Charmsaz, Yeonju Cho, James Leatherman, Melissa Lyman, Jacob Mitchell, Luciane T. Kagohara, Michael G. Goggins, Kelly J. Lafaro, Jin He, Christopher Shubert, William Burns, Lei Zheng, Elana J. Fertig, Elizabeth M. Jaffee, Richard A. Burkhart, Won Jin Ho, Jacquelyn W. Zimmerman

**Affiliations:** Department of Oncology, Sidney Kimmel Comprehensive Cancer Center, Johns Hopkins University School of Medicine, Baltimore, Maryland; Convergence Institute, Johns Hopkins University School of Medicine, Baltimore, Maryland; Bloomberg∼Kimmel Institute for Cancer Immunotherapy, Johns Hopkins University School of Medicine, Baltimore, Maryland; Department of Pathology, Johns Hopkins School of Medicine, Baltimore, Maryland; Department of Surgery, Johns Hopkins University School of Medicine, Baltimore, Maryland; Department of Genetic Medicine, Johns Hopkins School of Medicine, Baltimore, Maryland; Department of Biomedical Engineering, Johns Hopkins School of Medicine, Baltimore, Maryland; Department of Applied Mathematics and Statistics, Whiting School of Engineering, Johns Hopkins University, Baltimore, Maryland; Institute for Genome Sciences, Department of Medicine, and Greenbaum Comprehensive Cancer Center, University of Maryland School of Medicine, Baltimore, Maryland

## Abstract

Pancreatic ductal adenocarcinoma (PDAC) carries an extremely poor prognosis, in part resulting from cellular heterogeneity that supports overall tumorigenicity. Cancer associated fibroblasts (CAF) are key determinants of PDAC biology and response to systemic therapy. While CAF subtypes have been defined, the effects of patient-specific CAF heterogeneity and plasticity on tumor cell behavior remain unclear. Here, multi-omics was used to characterize the tumor microenvironment (TME) in tumors from patients undergoing curative-intent surgery for PDAC. In these same patients, matched tumor organoid and CAF lines were established to functionally validate the impact of CAFs on the tumor cells. CAFs were found to drive epithelial-mesenchymal transition (EMT) and a switch in tumor cell classificiaton from classical to basal subtype. Furthermore, we identified CAF-specific interleukin 8 (IL-8) as an important modulator of tumor cell subtype. Finally, we defined neighborhood relationships between tumor cell and T cell subsets.

**Statement of Significance:** This multidimensional analysis highlights the diverse role CAFs have in influencing other cell types in the TME, including epithelial-derived tumor and infiltrating immune cells. Our methods provide a platform for evaluating emerging therapeutic approaches and for studying mechanisms that dictate tumor behavior in a manner that reflects patient-specific heterogeneity.

## Introduction

Pancreatic ductal adenocarcinoma (PDAC) is a debilitating disease with a dismal five–year survival rate of 13% ^1^. These poor outcomes are due to numerous factors, including late–stage diagnosis, resistance to current therapeutic regimens, and importantly, a complex heterogenous tumor microenvironment (TME)^2–5^. The TME consists of a diverse and evolving ecosystem of epithelial, endothelial, immune cells, and cancer associated fibroblasts (CAFs)^6–8^. CAFs are present in abundance in the PDAC TME and have been implicated as drivers of an immunosuppressive milieu that provides a niche for tumor cells, promotes tumor growth and ultimately supports metastasis^6,9–14^.

Molecular and cellular classification, including both CAF and tumor cell subtypes, may influence clinical decision making with both current and emerging therapeutic approaches^15^. Two main subtypes of tumor cells have been described; a classical subtype that retains epithelial markers and is often well-differentiated, and a basal subtype that is quasi-mesenchymal with poorer responses to chemotherapy and worse overall survival^16^. Additionally, CAF subtyping has associated unique gene expression programs with specific roles within the TME. The most common CAF classifications include inflammatory (iCAF) with the capacity to secrete IL-6, a myofibroblastic (myCAF) subtype defined by high levels of alpha smooth muscle actin (αSMA) expression, and an antigen presenting (apCAF) subtype defined by expression of HLA-DR and antigen processing and presentation capacity^14,15,17–20^. While these distinctions underscore TME complexity in PDAC, the dynamic and nuanced interactions between CAFs and epithelial tumor cells remain incompletely understood.

In an effort to start to untangle these relationships, we used multi-omics platforms (transcriptomics and spatial proteomics) to examine how CAFs alter tumor biology through signaling mechanisms within the PDAC TME. We developed an approach to integrate multiplex imaging data from patient tissue slides with functional *in vitro* study of patient-derived organoid (PDO) – CAF cocultures. This approach demonstrated a spatial relationship between iCAFs to classical tumor cells and myCAFs to basal tumor cells. These data suggest that CAF proximity to tumor cells can promote a classical to basal phenotype switch and drive epithelial-mesenchymal transition (EMT) signaling. This switch was associated with decreased CAF-derived IL-8 secretion and impaired maintenance of a classical tumor cell phenotype amongst the tumor epithelial cells. Additionally, tumor regions with increased basal gene expression were associated with increased infiltration of activated T cells, suggesting that CAF-epithelial cross-talk serves to define the immune landscape in this disease. Thus, this study provides new data supporting CAF signaling in shaping the cellular and behavioral heterogeneity in the PDAC TME. These findings can be used to explore rational approaches to improve therapies for this difficult-to-treat disease.

## Results

### Spatial proteomics of patient tumors defines distinct CAF and tumor cell neighborhoods

Two subtypes of tumor cells (classical and basal) and three subtypes of CAFs (iCAF, myCAF, and apCAF) are accepted as important contributors to PDAC heterogeneity^16,18^. To further delineate these subtypes in a patient-specific manner, we comprehensively analyzed tumor specimens from 15 patients with PDAC who underwent surgical resection at our institution (Table 1). Imaging mass cytometry (IMC) was utilized to label human tissue with a panel of 43 antibodies (Supplemental Table 1) designed to capture spatially resolved profiles of CAFs, epithelial cells, and immune cells. Specific regions of interest (ROI) were selected for annotation by an expert pathologist (J.W.L.) (Supplemental Figure 1). To ensure similar sampling of the TME from every patient, ROIs were selected for malignant epithelial rich or stroma rich areas, and then further divided as immune rich or immune poor (Figure 1A). To annotate immune rich or poor regions, dual immunohistochemistry staining for CD3 (T cells) and CD68 (myeloid cells) was performed on sequential slides and a grid system was applied to account for size of ROI to be ablated for IMC (Supplemental Figure 1). We intentionally avoided tertiary lymphoid structures (TLS) when selecting ROIs, as these regions have been shown to be restricted to tumor edges^21^. The IMC panel was curated to ensure comprehensive analysis of CAF subtypes as well as endothelial, epithelial, and immune cell subtypes (NK, B, myeloid, and multiple subsets of T cells)(Figure 1B). All cell types were analyzed by relative expression of each marker included in the panel (Figure 1C). Clustering analysis identified groups of epithelial cells (CK+), CAFs (αSMA^+^, VIM^+^, PDGFR^+^, and/or FAP^+^, and CK^lo^), immune cells (both myeloid [CD68+] and lymphoid [CD3+ and CD20+]) as the most highly identified clusters (Figure 1C). The total number of cells and distribution of cell types acquired for each patient reflected patient-specific heterogeneity, and importantly, all of the broad cell types were represented in every patient sample (Figure 1D). Proportionally, CAFs (αSMA^+^, VIM^+^) and tumor cells (basal [KRT17^+^] or classical [TFF1^+^]) were the most prevalent cell types across our heterogenous patient cohort (Figure 1D-E).

**Figure 1.**
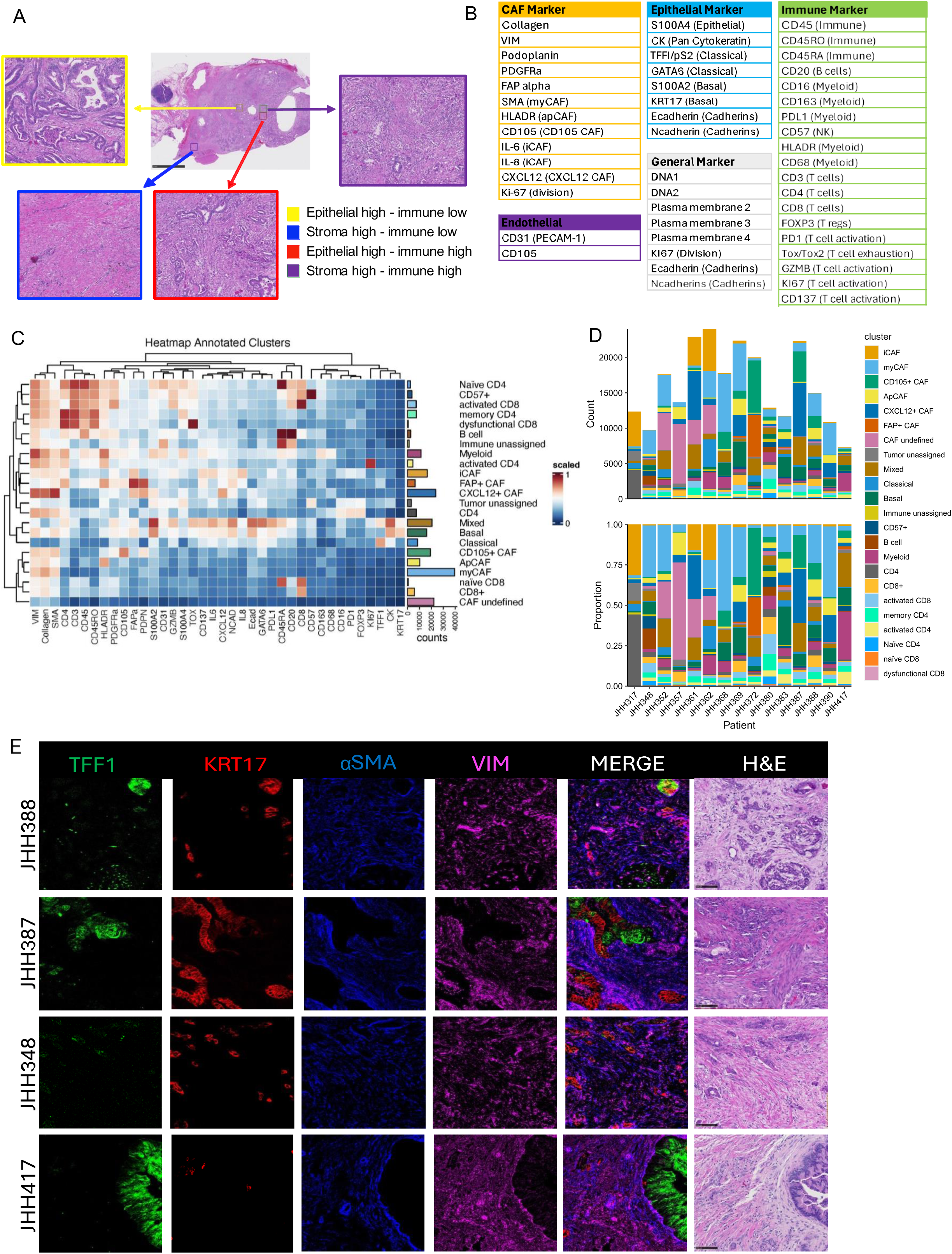
Spatial proteomic profiling of patient tumors identifies defined clusters and regions of cells in PDAC. (A) Visualization of different regions that were acquired for IMC analysis - Stromal high (blue and purple), Tumor high (red and yellow) regions annotated by an expert pathologist (JWL). Immune high (red and purple) and immune low (blue and yellow) regions were chosen when possible. (B) IMC panel markers grouped by cell type. (C) Cell type heatmap annotations by examining relative expression of relevant markers included in the IMC panel. (D) Total number of cells analyzed by region visualized as distribution by patient. (E) Representative images of 4 PDAC patients demonstrating classical markers (TFF1 - Green), basal markers (KRT17 - Red), myCAF (αSMA - blue), activated CAF (VIM – pink), Merge, and matching H&E images observed at 10x magnification (scale bar = 100µm), (N=15).

**Table 1.** Patient Demographics.

| Patient* | Age | Sex | Time to recurrence (mo) | KRAS NGS | Neoadjuvant treatment | Adjuvant treatment | Surgical Procedure<sup>#</sup> | Pathologic Stage |
| --- | --- | --- | --- | --- | --- | --- | --- | --- |
| JHH317 | 72 | F | 20.2 | G12D | Gemcitabine nab-paclitaxel and immunotherapy | Gemcitabine nab-paclitaxel and immunotherapy | DPS | T3N2 |
| JHH348 | 84 | F | 9.9 | G12R | Untreated | None | Whipple | T3N2 |
| JHH352 | 57 | F | 14.7 | G12V | FOLFIRINOX | FOLFIRINOX | Whipple | yT1N2 |
| JHH357 | 62 | M | 22.9 | G12V | FOLFIRINOX | Gemcitabine Capecitabine | Whipple | yT2N2 |
| JHH361 | 63 | M | 20.7 | G12D | FOLFIRINOX | Gemcitabine Capecitabine | DPS | yT2N0 |
| JHH362 | 83 | F | - | - | Untreated | - | DPS | T2N1 |
| JHH368 | 79 | F | 23.3 | G12R | Gemcitabine nab-paclitaxel | Gemcitabine | DPS | yT2N2 |
| JHH369 | 57 | M | 4.7 | G12D | FOLFIRINOX | Gemcitabine nab-paclitaxel | Whipple | yT2N1 |
| JHH372 | 57 | M | 7.0 | Q61H | FOLFIRINOX | Gemcitabine nab-paclitaxel | Whipple | yT2N0 |
| JHH380 | 85 | F | 6.2 | G12V | Untreated | None | DPS | T3 |
| JHH383 | 57 | F | 12.2 | G12R | Immunotherapy | FOLFIRINOX and Gemcitabine nab-paclitaxel | Total pancreatectomy and splenectomy | yT3N2 |
| JHH387 | 55 | M | 8.2 | G12D | FOLFIRINOX | - | DPS | yT4N2 |
| JHH388 | 52 | M | 5.9 | G12D | FOLFIRINOX | Gemcitabine nab-paclitaxel | DPS | yT3N0 |
| JHH390 | 54 | M | - | G12D | FOLFIRINOX | FOLFIRINOX Olaparib<sup>\$</sup> | Whipple | T2N0 |
| JHH417 | 72 | F | 9.8 | G12V | Untreated | FOLFIRINOX | Whipple | T2N2 |

\*JHH348, JHH362, JHH380 were included in pilot bulk RNA sequencing; JHH388 and JHH417 were added to the IMC cohort and are not represented in the RNA sequencing data; JHH390 cocultured organoid sample did not meet quality parameters for analysis and thus is omitted from direct comparisons between monoculture and cocultured organoids

With the primary goal of directly evaluating the relationship between CAFs and tumor cells, we clustered our data to look exclusively at CAF and tumor cell populations, both across the cohort and at the level of individual patients (Figure 2A-B). We performed nearest neighbor analysis to understand interactions between CAF and tumor cell subsets (basal, classical, or mixed). Mixed tumor cell designation was assigned to tumor cells expressing both basal and classical markers with these data suggesting plasticity of the epithelial cohort between subsets and a transitional state. Upon enumerating the top 3 nearest cells within a 4-μm distance radius from each of the basal, classical, or mixed tumor cells and filtering the counts by CAF subtypes only, we found that iCAFs were the most frequent cell type nearest in distance to classical tumor cells while myCAFs were most frequently nearest to basal cells (Figure 2C). apCAFs, which represented a smaller minority of CAFs (Figure 2D), were also observed frequently as a nearest neighbor of basal cells (Figure 2C). These data suggest CAF and epithelial subtypes are paired together uniquely in spatial neighborhoods (iCAF/classical and myCAF/basal) across a more broadly heterogenous TME (Figure 2D). These associations were also visually confirmed within ROIs displaying both classical-rich and basal-rich areas of tumor cells (Figure 2E). Taken together, these data demonstrate a distinct spatial coordination among CAF and tumor cell subtypes, suggesting CAF subtypes are influencing the phenotypes of nearby tumor cells.

**Figure 2.**
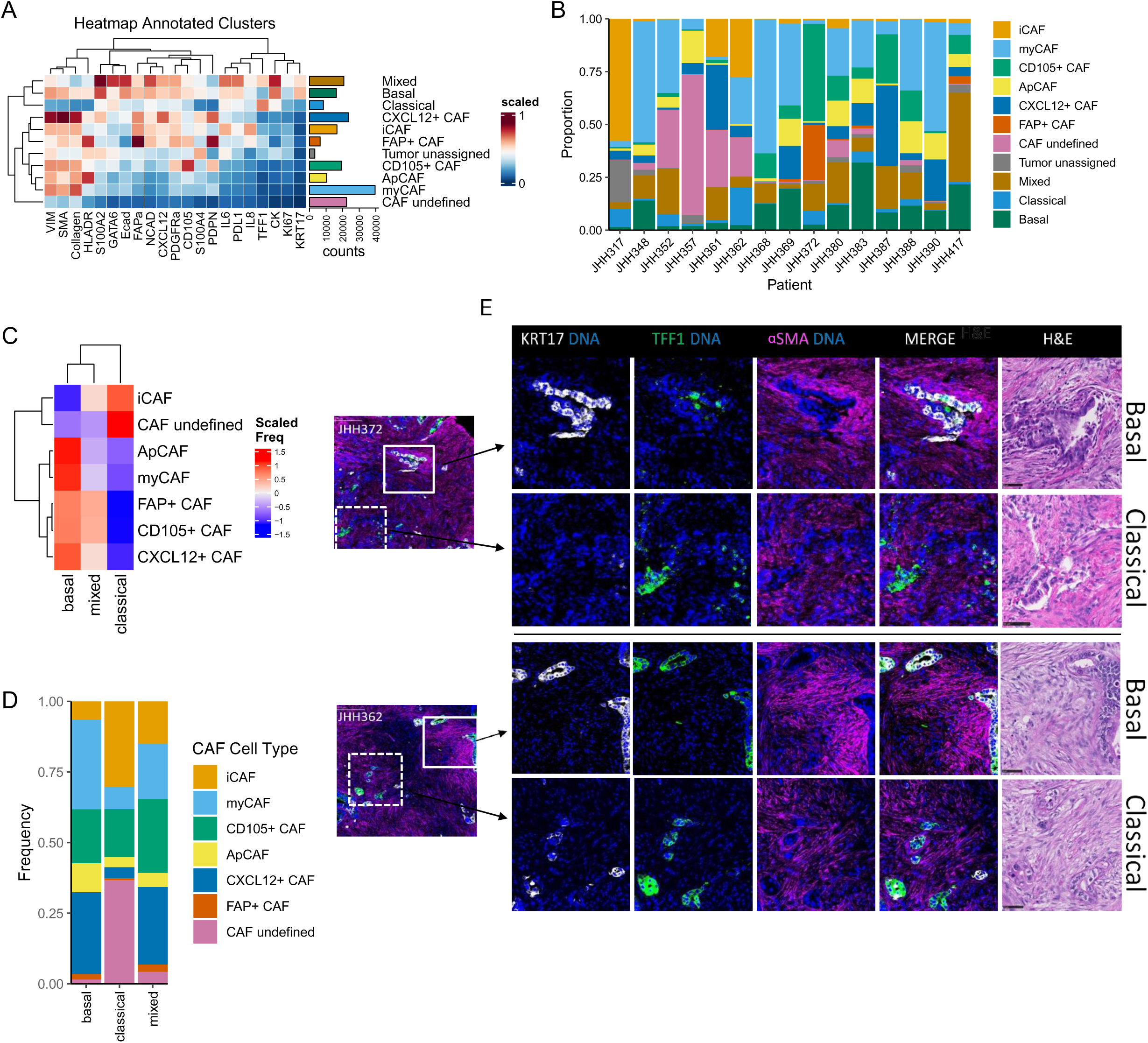
Spatial proteomics analysis reveals myCAFs are spatially located nearest to basal PDAC tumor cells. (A) Cell type heatmap for data subset for CAFs and tumor cells. Cell type annotations were determined by relative expression of relevant markers included in the IMC panel. (B) Total number of cells analyzed by region with visualization of distribution by patient. (C) Heat map showing nearest neighbor analysis identifying distinct CAF subtypes that are closest to basal, classical, and mixed tumor cells. (D) Stacked barplot showing cell type frequencies of CAF subtypes that are present nearest to basal, classical, and mixed tumor cells dictating increased myCAF frequency near basal cells, and increased iCAF frequency near classical cells. (E) Representative images of IMC and H&E for 2 patients that have basal rich and classical rich regions displayed in one ROI. Images depict more myCAF presence (αSMA+) and intensity in basal rich (KRT17+) regions (top row for each patient), H&E scale bar = 50µm.

### Bulk RNAseq of patient-matched PDO and CAF cocultures define CAFs as drivers of EMT in PDOs

To more comprehensively evaluate CAF-tumor cell crosstalk within the IMC defined neighborhoods, we completed bulk RNA sequencing of flow-sorted CAF and PDO cocultures from 12 patients with PDAC. Generalizability of these data to diverse PDAC patient cohorts were preserved given that the current set of samples represents 3 untreated patients, 6 previously treated with neoadjuvant FOLFIRINOX (5-FU, irinotecan, oxaliplatin, leucovorin), and 3 having received gemcitabine and other neoadjuvant treatments (Figure 3A-3B, Table 1). Data were first evaluated by computing principal components, which demonstrated that cell type was the predominate source of variation in the data and distinguish CAFs from organoids (Figure 3C). Further visualization of coavariates on the principal component analysis did not find significant variation in the transcriptional data related either to culture conditions (coculture or monoculture as control, Figure 3D) or individual patients (Figure 3E). While not a dominant source of variation at a global transcriptional level in the principal component analysis, we still hypothesized that the co-culture would impact the transcriptional profiles relative to monoculture. Differential analysis of gene expression compared cocultured PDOs to monoculture PDOs to identify CAF-mediated effects (Figure 3F). Following PDO-CAF coculture, there was a significant increase in the expression of genes associated with fibrogenesis and malignant progression, including *POSTN, DCN, COL1A2,* and *CHD1* and *MMP2*, respectively (Figure 3F). We further identified enrichment of epithelial-mesenchymal transition (EMT), TNFα response, E2F targets, KRAS and MYC targets in in coculture PDOs relative to monoculture using the Hallmark gene sets from the Molecular Signatures Database^22^ (Figure 3G).

**Figure 3.**
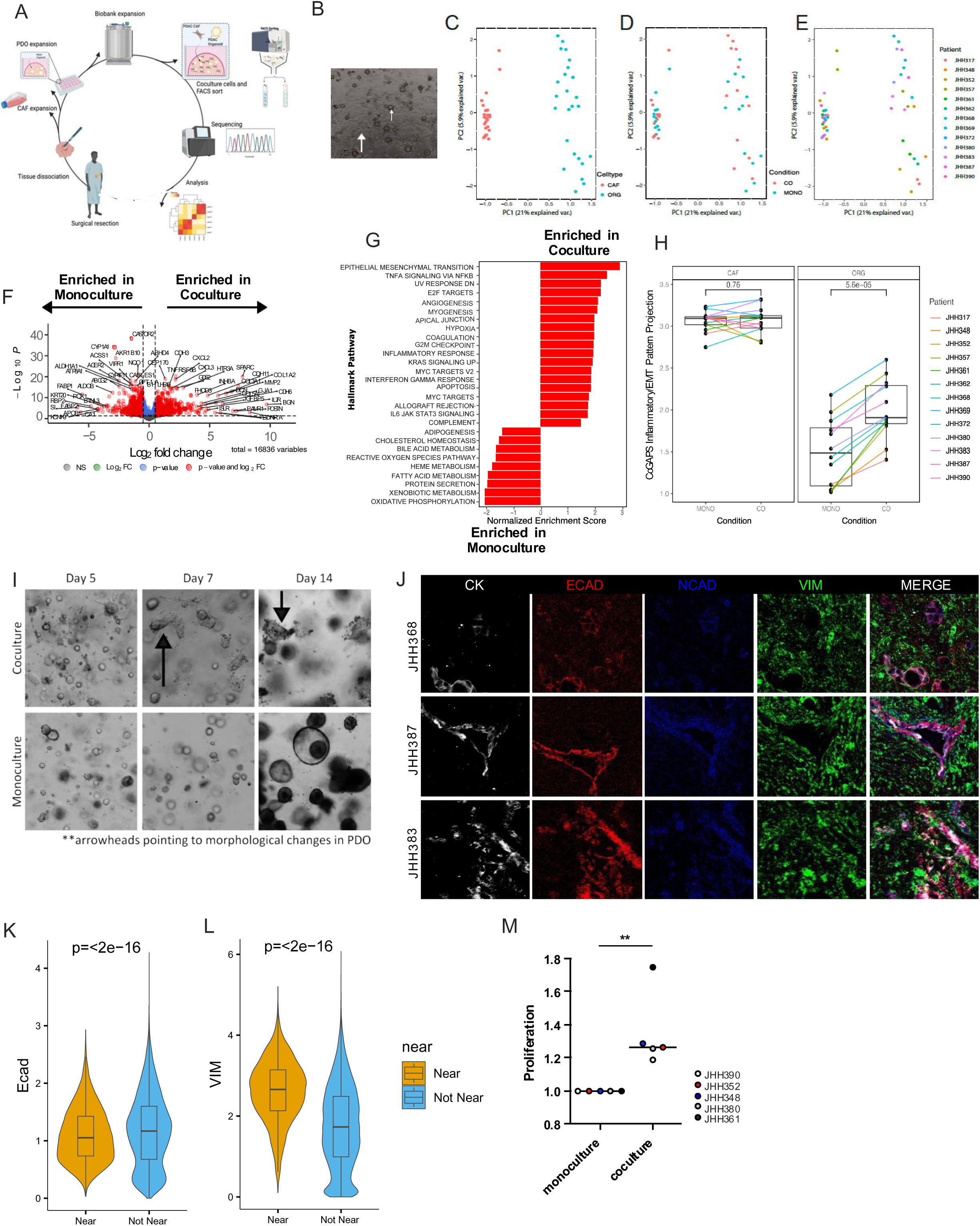
RNA sequencing of patient matched PDO and CAF coculture reveals that CAF presence drives heightened EMT in PDOs. (A) Graphical schematic depicting creation of the biobank from patient tumors and workflow for performing RNA sequencing. (B) Representative image of CAF – PDO coculture prior to harvest for RNA sequencing. Large arrow points towards CAF and smaller arrow towards PDO. (C) Unsupervised clustering of viable cells from 12 patient CAF – PDO cocultures represented as PCA plot. (D) Resolution of coculture and monoculture cells represented by PCA plot. (E) Resolution of each patient represented by PCA plot. JHH 390 co-org did not meet quality parameters upon sequencing, so it was excluded from analysis. (F) Differential gene expression of PDO samples that are monocultured (left) compared to cocultured (right). (G) Overrepresented MSigDB hallmark gene sets in PDO samples that are monocultured (left) compared to cocultured (right). (H) ProjectR transfer learning of gene signatures inferred in single-cell RNA-seq data of PDAC tumors (*35*) onto the RNA sequencing data demonstrates that the pattern associated with inflammatory signaling and EMT inferred in our prior study is enhanced in only PDO cells from coculture relative to PDO cells from monoculture in this study, p=5.6e-5 by two-tailed paired students T-test. (I) Representative time-course images of monoculture PDO and coculture CAF - PDO demonstrating the morphologic changes that occur in PDO over the course of two weeks that is not seen in the monoculture PDOs, images taken at 10X on Echo Rebel microscope. (J) Representative images from IMC showing traditional EMT markers, E-cadherin, N-cadherin, and Vimentin. Quantification of E-cadherin (K) and vimentin (L) expressed in basal tumor cells near a CAF or not near a CAF. (M) Proliferation of tumor cells in monoculture setting or coculture setting. 5 patients were profiled with 4 technical replicates per patient. Significance is measured as: ****, p<0.0001; ***, p<0.001; **, p<0.01; *, p<0.05; ns, not significant.

These findings of enhanced EMT in coculture support our prior work examining the global transcriptional impact of CAFs on tumor cells^23^. In this study, analysis of a comprehensive atlas of public-domain single-cell RNA sequencing data of PDAC tumors identified a gene expression pattern in tumor cells associated with elevated inflammatory, fibrogenic, and EMT pathway activity^23^. We referred to this as the “Inflammatory/EMT” pattern, and enrichment of this pattern appeared to be driven by the presence of, and regulation by, CAF signaling that was observed in co-culture data of PDOs from three treatement-naïve patients. In the full 12 patient cohort explored here, we identified CAF-PDO coculture as inducing increased expression of Inflammatory/EMT gene pathways, as compared to those from monoculture (Figure 3H). This suggests that CAFs signaling to epithelial tumor cells plays a critical role in the promotion of tumor inflammation, fibrogenesis and EMT. In addition to the broader transcriptomic findings, we observed morphologic changes consistent with EMT in cocultured PDOs compared to monoculture counterparts (Figure 3I). Further evidence to support CAF-directed EMT was found by directly evaluating patient tissues by IMC-staining for expression of E-cadherin, N-cadherin, and vimentin (Figure 3J). We hypothesized tumor cells in close proximity to CAFs would demonstrate increased evidence of EMT through loss of E-cadherin and increased vimentin. Based on IMC analysis, E-cadherin expression was in fact greater in tumor cells distant from a nearby CAF (Figure 3K), while vimentin expression was greater in tumor cells near a CAF (Figure 3L). There was limited difference in N-cadherin expression amongst tumor cells evaluated. Our previous study showed that at a single-cell level induction of EMT from fibroblasts is mutually-exclusive with proliferation in the neoplastic cells^23^.

Nonetheless, prior work suggests CAFs may also promote a proliferative phenotype in tumor epithelial cells^24^. We therefore examined PDO proliferative capacity in coculture by adapting a luciferase transduction protocol to accommodate PDOs. In our coculture there was increased proliferation of the PDOs, as compared to monoculture PDO controls (Figure 3M). Taken together, these data support the hypothesis that CAFs are critical to and controlling drivers of EMT and proliferation in PDAC tumor cells at a population level.

### CAFs promote changes in PDO gene expression from classical to basal tumor cell classification

We next returned to the classical/basal tumor cell classification system, first described by Moffitt and colleagues, to examine gene signatures in our PDOs and the tumor-specific heterogeneity in the capacity of CAFs to alter tumor cell classification^24,25^. In PDO monoculture there was heterogeneity in relative expression of key classical and basal classifier genes (Figure 4A). As CAFs were introduced into cultures, PDO gene expression was often altered and profiling revealed plasticity of gene expression in the epithelial compartment in a manner dependent on the presence of CAFs (Figure 4B). In contrast to the increase of EMT in the Inflammatory/EMT pattern defined from our atlas, transition to a more basal-like phenotype was only observed in a subset of the organoids. In the 5 patients for whom the gene expression was most dynamic, there was a significant shift toward increased basal gene expression (Figure 4C). There were no patient samples in which tumor cell exposure to CAFs through coculture induced a more classical gene expression pattern. Morover, this state transition appeared independent of the initial subtype classification of the organoid monoculture, although the 4 lines with the highest basal scores in monoculture did not significantly change subtype.

**Figure 4.**
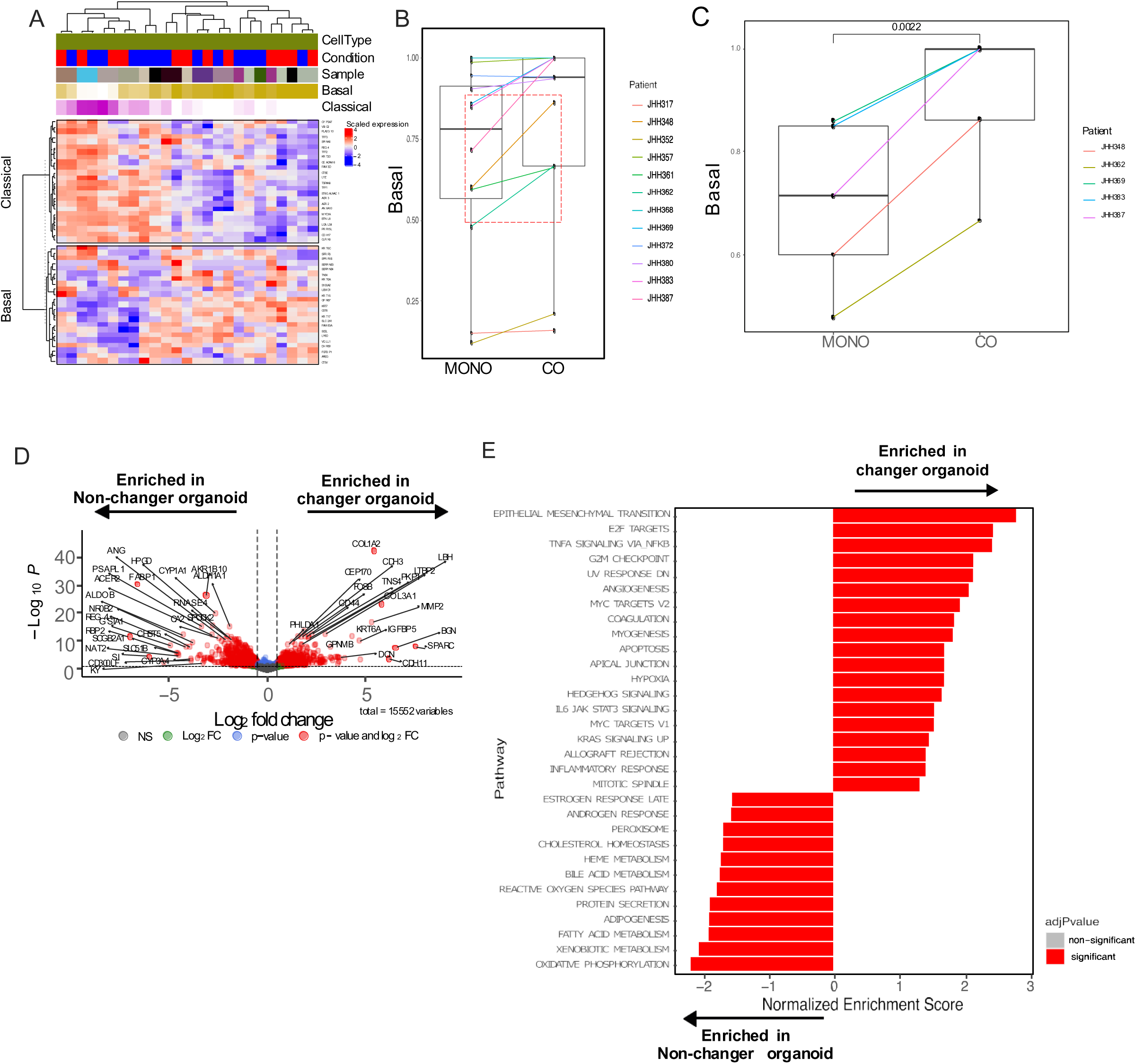
CAFs drive changes in PDO gene expression from classical to basal tumor cell classification. (A) Heatmap displaying basal and classical gene expression in PDOs between coculture and monoculture samples. (B) Basal gene expression on a per patient basis. Red hashed box demonstrates the PDOs that upregulate basal gene expression in coculture PDOs (n=5) (p=0.0022). (C) Top 5 PDO lines that change from classical to basal in coculture setting. Comparisons of conditions are statistically supported using the two-tailed students t-test with equal variance p=0.0022. (D) Differential gene expression showing the genes upregulated in n=7 lines that do not shift (left) and genes upregulated in PDO that become basal n=5 (right) visualized by volcano plot. (E) Overrepresented MSigDB Hallmark gene sets in PDO samples that do not shift (left) compared to PDO samples that shift – “changers” (right).

A quantitative assessment of gene expression signatures identified a strong basal shift in these 5 patients, with increased relative basal expression of global gene programs by at least 14% and up to 28% (Figure 4C). Assessment of the expression of specific basal genes such as *KRT6A, COL1A2*, and cadherins, as well as cancer stem cell genes such as *CD44,* demonstrated increased pathway signaling with CAF coculture suggesting CAFs are driving a more basal and more aggressive PDO phenotype in part through these critical genes (Figure 4D). In these samples, and in keeping with a more basal phenotype, increased PDO tumorgenicity was inferred by Hallmark pathway analysis, showing upregulation of EMT and proliferation by E2F targets, as well as upregulation of angiogenesis and hypoxia signaling pathways (Figure 4E). To validate our coculture findings as relevant to patient disease, we returned to primary patient tissues and used multi-plex RNAscope to probe for basal (*KRT17* and *KRT6a*) and classical (*TFF1* and *GATA6*) RNA transcripts (Supplemental Figure 2A) on tissue sections. Similar to our coculture findings, there was a range of expression of representative basal and classical genes across patients at baseline. Additionally, there was a direct correlation between the selected basal RNAs and classical RNAs reinforcing their co-expression and validity as markers to designate tumor cell subtype (Supplemental Figure 2B). To determine if the basal phenotype was associated with an abundance of CAFs as seen in our coculture, we used trichrome staining to infer CAF density also at a whole-tissue scale. In samples demonstrating heavy trichrome staining, and therefore a high CAF presence, the patient-specific PDO subtype was more likely to be either strongly basal at baseline or to be a PDO that could be induced towards a basal phenotype with CAF coculture (Supplemental Figure 3). These data further reinforce the hypothesis that dynamic CAF-driven signaling helps to regulate epithelial gene expression that is reflected in clinically-impactful tumor cell classification.

We then sought to determine whether co-culture with PDOs also led to changes within the CAFs (mean myCAF signature of approximately 0.985 vs. 0.975, respectively; Supplemental Figure 4A-B). We hypothesized that this could be due to the limitations of bulk RNA sequencing and, therefore, queried specific CAF markers with qPCR to examine gene expression from cocultures relative to monocultures. Using this more granular approach, we identified significant changes in expression of *ACTA2* (αSMA), *VIM* (vimentin), and *COL1A1* suggesting increased myCAF gene programming in coculture (Supplemental Figure 4C).

### CAFs drive basal gene expression in PDOs through secreted proteins

While our data suggest proximity between CAFs and tumor cells alters tumor cell classification and drives tumor EMT, it remained unclear if this epithelial cell plasticity was due to direct cell-cell interaction or a result of signaling mediated by the CAF secretome. To evaluate the role of CAF-secreted factors on tumor cell molecular states, we treated PDOs with CAF-conditioned media and queried classical and basal gene expression changes using qPCR. Overall, culturing in CAF conditioned media induced altered PDO gene expression from classical towards increased basal programming (Figure 5A). In exploring the functional implications of CAF-conditioned media, and to examine the capacity of the secretome to alter cell migration, we performed a wound healing assay using Panc 10.05 cells^25^. Cells treated with CAF conditioned media from basal patient lines demonstrated an increased rate of wound closure as compared to those cultured with conditioned media from CAFs matched to pateints with classical PDO lines (Supplemental Figure 5A-B), suggestive of a more basal-like behavior.

**Figure 5.**
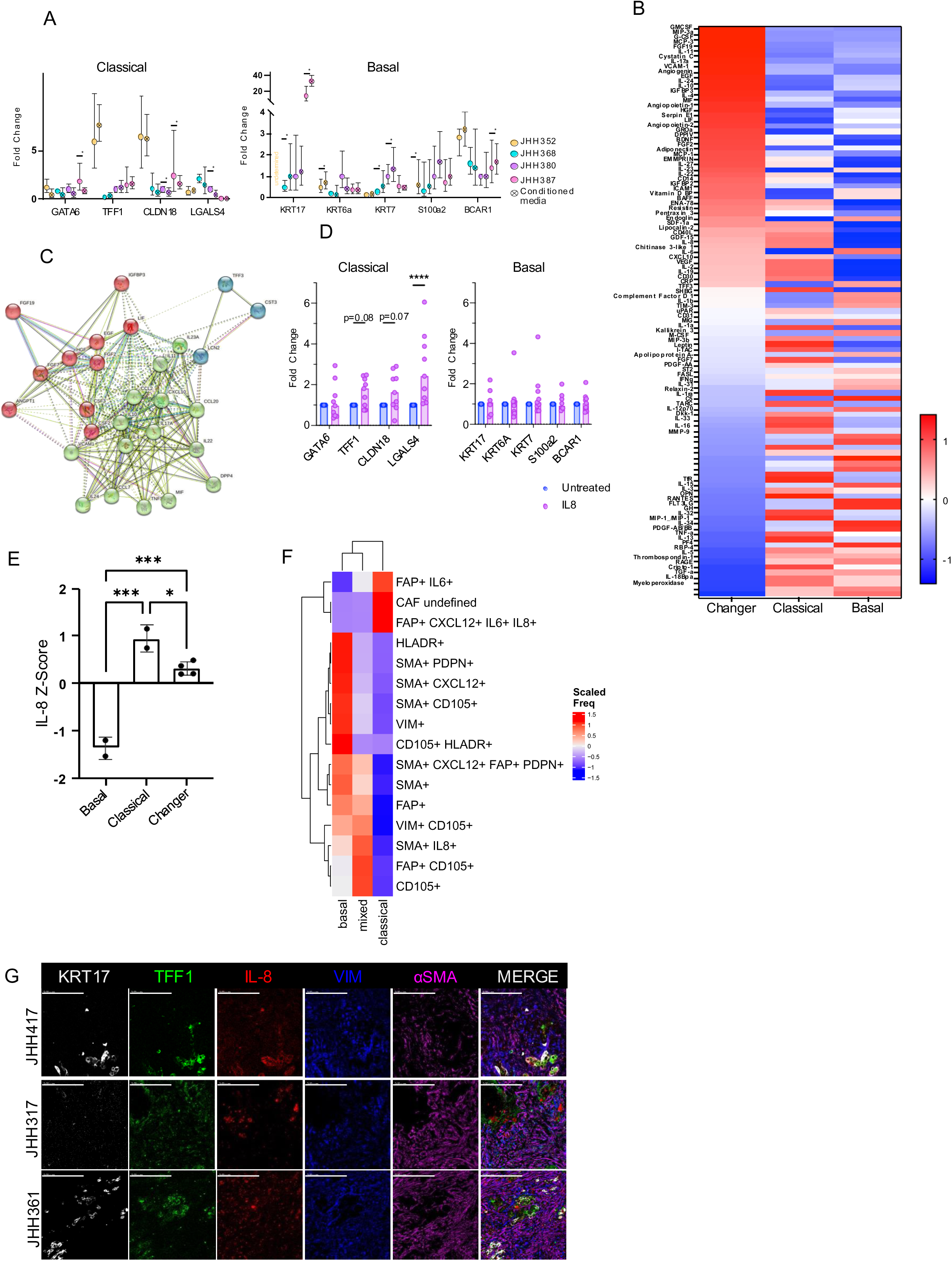
CAFs that drive basal gene expression in PDO secrete distinct proteins such as IL-8. (A) CAF CM treated PDOs have decreased classical gene expression (left) by qPCR for GATA6, TFF1, CLDN18, and LGALS4. CAF CM treated PDOs show increased basal gene expression (right) by qPCR for KRT17, KRT6a, KRT7, S100a2, and BCAR1. JHH352 did not amplify for KRT17. (B) Heatmap of conditioned media secretome analysis from CAFs that change PDO from classical to basal (far left), CAFs that do not change classical, CAFs that do not change basal (far right). Results show normalized Z-Score of biological replicates. (C) StringDB protein interaction network. (D) Classical phenotype gene qPCR readout from PDOs treated with rIL-8 compared to untreated (left), basal phenotype gene qPCR readout from PDOs treated with rIL-8 compared to untreated (right). (E) IL-8 secretion from proteome screen (B) analyzed by normalized Z score. Comparisons of rIL-8 and untreated conditions are statistically supported using the two-tailed students t-test with equal variance in PRISM (V9.2.0 [283]). (F) Resolved heatmap of CAF defined markers where IL-8+ CAF are nearest to classical or mixed tumor cells. (G) Representative images IMC of 3 patients showing colocalization of IL-8 and classical tumor markers. Significance is measured as: ****, p<0.0001; ***, p<0.001; **, p<0.01; *, p<0.05; ns, not significant.

To identify secreted proteins that may be responsible for these classical vs. basal molecular and functional differences, we analyzed the secretome of conditioned media from patient-derived CAFs, comparing media from 3 CAF lines: CAFs from a line that maintains a classical PDO, CAFs from a line that maintains a basal PDO, and CAFs that drive classical-to-basal shift in their matched corresponding PDO. Upon profiling the CAF secretomes, we found 49 secreted proteins unique to patients that change from classical to basal gene expression (Figure 5B). We input these data into String DB to infer specific networks of cell crosstalk driven by protein interactions and selected 3 dominant clusters^26^. These included immune related proteins, growth factor related proteins, and cell mobility and protein-protein interaction proteins (Figure 5C). Further pathway-specific analysis suggested that 14 proteins were significantly enriched in this patient group compared to other groups. However, while these results infer proteins distinctly enriched in the group with plasticity, they do not indicate whether the proteins favor classical or basal. To determine the functional roles of these enriched proteins, we treated PDOs with recombinant proteins for these 14 hits to evaluate their capacity to alter PDO gene programming from a classical to basal phenotype (Supplemental Figure 5C-D). Of the 14 proteins screened, we identified IL-8 as a key protein in the secretome that maintains gene expression corresponding to a classical phenotype in tumor cells (Figure 5D). Yet, IL-8 exposure resulted in no meaningful change to the epithelial cell gene expression of basal targets *KRT17* and *S100A2* (Figure 5D). Thus, it appears that CAF-derived IL-8 signaling in classical phenotype tumors, when lost, can induce basal gene expression. When we went back to our screening data to compare IL-8 secretion across CAF lines, IL-8 secretion was the highest in the CAF line that was associated with classical epithelial gene expression, diminished in the CAFs that induced basal gene expression in their corresponding PDO, and nearly undetectable in the basal CAF line (Figure 5E). This suggests that CAF secreted IL-8 is required to keep tumor cells in a classical tumor designation when evaluated within the context of a competent 3D model of the TME. Upon transition from our reductionist coculture to patient tissue sections, we found that FAP^+^, CXCL12^+^, IL6^+^, IL-8^+^ CAFs were nearest to classical tumor and mixed tumors (Figure 5F-5G). These data show that IL-8 secreted by CAFs promotes preservation of the classical tumor subtype and identify an additional avenue of regulatory function for IL-8 within the TME.

### Activated T cells are associated with basal tumors

Finally, since the CAF secretome analysis identified differences in expression of immunomodulatory ligands, we broadly examined the relationship between immune cell type in the TME and the heterogeneity in myCAF/basal and iCAF/classical neighborhoods. There has been limited clinical success employing immune checkpoint blockade to treat PDAC^27^. The ineffectiveness of immune checkpoint blockade has been attributed to immune evasion, low neoantigen burden, and immunoediting; recently, CAFs have been hypothesized to affect the immune landscape of the disease^28,29^. IMC analysis showed that among the immune cells, T cells were the nearest immune cells to all tumor cell subtypes (Figure 6A). Based on our results showing that IL-8, a cytokine involved in T cell suppression and resistance to checkpoint immunotherapy^30^, is an important determinant of classical phenotype, we hypothesized that the T cell frequency may be lower near classical cells. Indeed, T cells were seen at highest frequency near basal tumors, at intermediate levels near mixed tumor cells, and at lowest frequency near classical tumors (Figure 6B). Effector CD4 and CD8 T cells comprised the largest fractions of T cells, most noticeably in the basal and mixed neighborhoods (Figure 6B-C). Collectively, these data demonstrate that the character of the adaptive immune infiltration in the TME is heterogenous and associated with distinct tumor cell subtype. Future investigation examining immune recognition and function of those cells nearest to basal cells and the spatial coordination of immune cells nearest to classical cells could aid in better understanding cellular infiltration, recognition, and immunotherapy resistance in PDAC. Furthermore, exploring the direct impact of CAF-derived secreted factors including IL-8 on adaptive immune populations is warranted.

**Figure 6.**
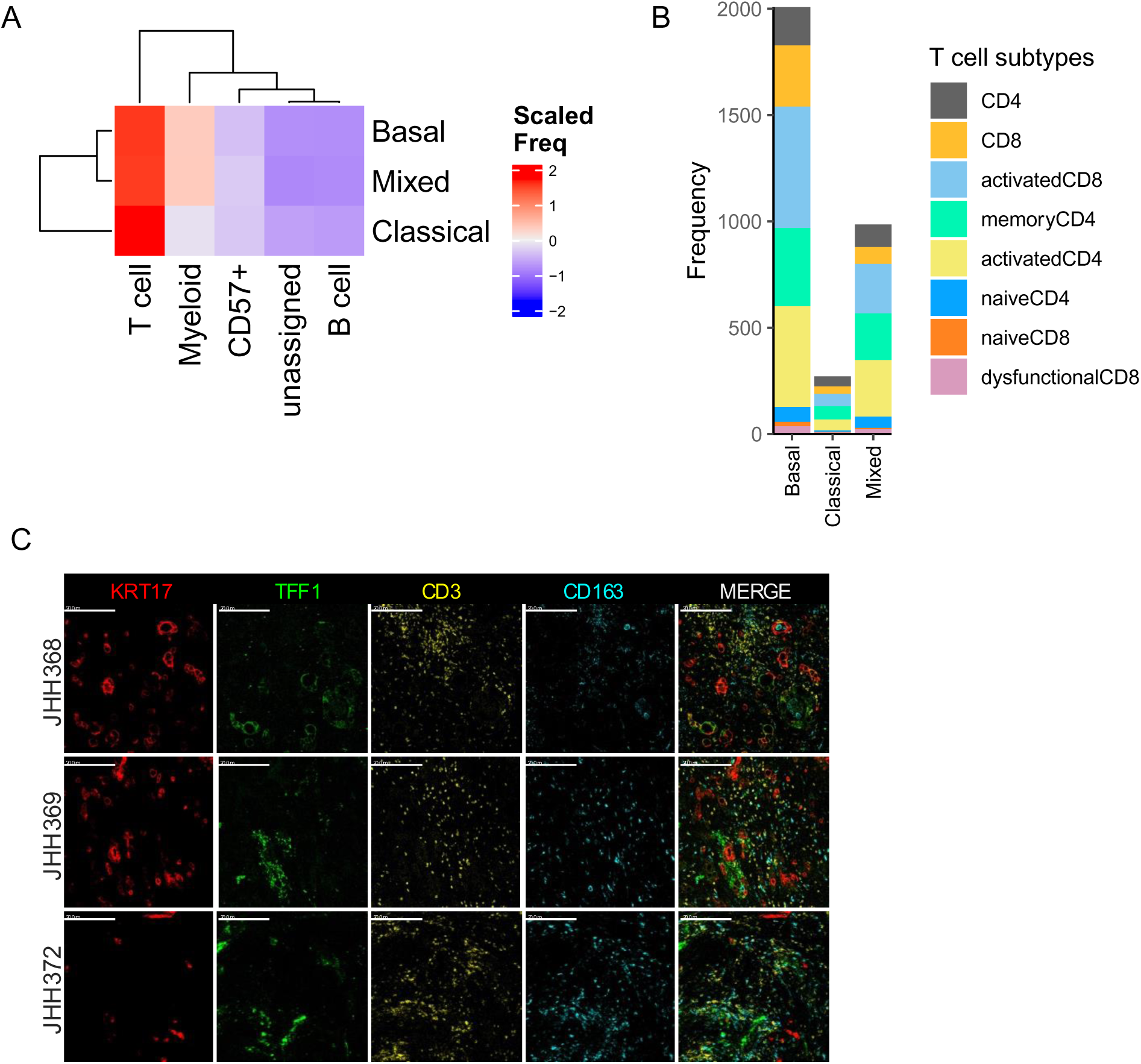
T cells are located in closest proximity to tumor cells. (A) Heatmap of nearest neighbor analysis of immune cells and tumor cells where T cells are the predominant immune cell near all classifications of tumor cells both my proximity (A) and by frequency (B). (C) Representative images of patient IMC showing tumor cells (KRT17+, TFF1+) and immune cells (CD3+, CD163+) colocalizing.

## Discussion

Here, we report a multidisciplinary study using comprehensive IMC to complement novel TME culture methods. We employed a living biobank of patient-matched CAFs and PDOs enabling us to broadly recapitulate the functional biology of CAFs *in vitro* and explore CAFs as critical regulators of the PDAC TME. By leveraging a combination of high dimensional data and patient-derived coculture, we identified CAF driven EMT, a neighborhood specific association of IL-8 secreting iCAF cells with classical tumor cell designation and loss of IL-8 expression associated with basal tumor designation. Finally, we identified T cells as the most abundant immune cell type associated with tumor-CAF neighborhods, with increased abundance in the basal cell and myCAF neighborhoods.

CAF promoted EMT is a complex phenomenon that can occur across tumor types. Prior work has shown that epithelial cells can lose expression of E-cadherin and transform into mesenchymal cells with a greater ability to migrate through the basement membrane, promote extracellular matrix deposition by stromal cells, and increase resistance to chemotherapeutics^31–33^. These malignant epithelial cells can then reach distant organs such as the liver or lung and establish metastatic deposits^34,35^. These data further support a therapeutic emphasis on reprogramming CAFs to limit EMT and therefore prevent metastatic spread. In this current study, we found that our transcriptional signature of CAF-induced EMT observed in previously human tumors^33,34^ is also associated with EMT transitions in PDO CAF co-culture independent of prior treatment or subtype. Additional studies were needed to identify specific mechanisms responsible for these pro-tumorigenic changes, and to distinguish EMT induction from subtype switching in PDAC. Here in our PDO model we identified one mechanism utilized specifically by CAFs to promote a transition from classical to basal tumor cell phenotype in a heterogenous manner with the capacity to influence patient-specific outcomes. We identified IL-8 as an important factor secreted by classical CAFs that acts to maintain the classical tumor cell phenotype. As the loss of CAF-derived IL-8 in the TME was associated with a basal tumor cell phenotype, and basal PDAC prognosis is particularly bleak, it is reasonable to explore the role of IL-8 maintenance in next-generation therapeutic studies.

The role of IL-8 in tumor behavior is complex and incompletely understood, particularly in PDAC. IL-8 has been implicated as an important chemokine in PDAC aggressiveness through *in vitro* work, with studies suggesting tumor cell expression of IL-8 as a hallmark of cells undergoing EMT^36^. However, there are fewer studies examining the impact of IL-8 that is secreted from stromal cells in the TME. Our findings suggest IL-8 is central to maintaining a classical tumor cell phenotype, implying that (1) IL-8 in the PDAC TME may have more of a functional duality than previously appreciated and (2) the source of IL-8 in the TME is not exclusively epithelial cancer cells. Consistent with our findings, Carpenter and colleagues previously described KRT17^high^CXCL8+ (an alternative name for IL-8) tumor cells as an intermediary phenotype between classical and basal and correlated with intratumoral myeloid abundance^37^. Unlike the work from Carpenter and colleagues, we exclusively examined the impact of IL-8 on tumor cell subtype. This is a critical caveat, as extensive literature supports an immuosuppressive role for IL-8 through the recruitment of neutrophils and other myeloid derived suppressor cells^38,39^.

Characterization of CAF heterogeneity remains challenging, given their diverse molecular functions and limitations of available model systems^9,18,40–42^. For example, CAFs have been implicated as having both tumor promoting and tumor restraining properties, furthering the challenges associated with classification^43,44^. These collective works demonstrate changes in stromal composition can have many effects on tumor development and phenotype. Additionally, PDAC tumor subtypes can be classified transcriptionally and can correlate with different types of stroma^16,45,46^. Our work, collectively with others, further reveals the heterogeneous landscape and spatial relevance of both tumor and non-epithelial cells in the environment, providing an additional source of gene expression, secreted proteins and other drivers of the dynamic behavior in the PDAC TME^19,47–51^.

Understanding CAF – tumor cell relationships at a single cell level is critical to more reliably represent the inherent intra-tumoral heterogeneity. This is particularly important when considering that many of the genes we have focused on are shared between CAFs and tumor cells, such as EMT markers like vimentin. While the results from this study further resolve the spatial relationships in the TME, there are also shortcomings to this study. Our IMC data are restricted to a panel that looks at these dynamic relationships broadly rather than explicitly in a single cell fashion. This presents a challenge when trying to relate cellular dynamics to cellular function more deeply and importantly, we were limited in the cell types we could comprehensively analyze. While myeloid cells are significant contributors to the TME, we were unable to fully characterize myeloid populations due to technical and antibody number limitations. Additionally, our CAF – PDO coculture system is an exemplary system to further study the TME. But, it is reductionist, and we did not explicitly explore immune interactions in these cocultures. Further immune phenotyping and therapeutic intervention opportunities warrant future exploration and provide an avenue for next steps integrating immune cells into this system.

Altogether, these findings demonstrate the importance of understanding the complex interplay between CAFs and tumor cells in the PDAC TME. We have defined a relationship whereby CAFs drive EMT and basal gene expression and also form distinct cellular neighborhoods of iCAFs/classical cells and myCAFs/basal cells. We also identified greater activated T cell presence within basal tumor regions, a finding that warrants additional exploration as we aim to improve immunotherapeutic strategies for PDAC. Future work will prioritize understanding the immunofunctional implications of these tumor cell and CAF relationships.

## Materials and methods

### Imaging Mass Cytometry (IMC)

Resected pancreas slides were baked at 60°C for 2 hours, dewaxed in histological grade xylene, then rehydrated in a descending alcohol gradient. Slides were incubated in Antigen Retrieval Agent pH 9 (Agilent,® S2367) at 96°C for 1 hour and blocked with 3% BSA in Maxpar® PBS at room temperature for 45 minutes. Immunohistochemical staining was done through individually conjugated mass cytometry antibodies. An antibody cocktail was prepared, detailed in Supplemental Table 1, and used to stain the slides at 4°C overnight. Custom antibodies were conjugated in-house, diluted to a concentration of 0.25 mg/mL to 0.5 mg/mL, then titrated empirically. Cell-ID™ Intercalator-Ir (Standard BioTools, 201192A) was diluted at 1:400 in Maxpar® PBS and used for DNA labeling. Ruthenium tetroxide 0.5% Aqueous Solution (Electron Microscopy Sciences, 20700-05) was diluted at 1:2000 in Maxpar® PBS and used as a counterstain. Images were acquired through the Hyperion Imaging System (Standard BioTools) at the Johns Hopkins Mass Cytometry Facility.

Images were prepared for analysis similarly to prior description^52^. In brief, images were segmented using nuclear (Ir191 and Ir193) and plasma membrane staining (IMC Segmentation Kit, Standard BioTools, TIS-00001). Sixty images were evaluated to assign pixel classifications and establish probability maps using *Ilastik*^53^. CellProfiler (v 4.2.4)^54,55^ was then used to generate segmentation masks for these images based on the resulting probability maps. The quality of segmentation was explored visually, and per-cell data were exported using *histoCAT*^56^. Clustering of individual cells was achieved using the relative expression of cell subtyping and functional markers using Phenograph^52^. Density of cell types was determined by dividing the number of cells detected *per* cluster by the area of tissue analyzed. Top neighbor analysis was performed by compiling the top 3 neighbors to each cell. Heatmaps were generated to display aggregated data and clearly label defined clusters. Representative images were prepared using MCD™ Viewer (Standard BioTools), overlaying multiple stains and adjusting the threshold to minimize background. These were then exported as 16-bit images. Box plots were generated in R v 3.6.3. using ggplot2.

### Patient sample acquisition and organoid and CAF line generation

Patients with PDAC undergoing surgical resection were enrolled in IRB-approved tissue acquisition protocols at Johns Hopkins Hospital, Table 1 (IRB: NA_00001584). PDOs were generated from patient surgical specimens following a combination of mechanical and enzymatic dissociation as previously described^57,58^. Organoid lines were maintained in Matrigel (Corning, 356234) with Human Complete Feeding Media (HCPLT media), detailed in Supplemental Table 2. For organoid passaging, media was aspirated and then Matrigel domes were resuspended in Cell Recovery Solution (Corning, 354253) and incubated on ice at 4°C for 45 minutes to allow Matrigel depolymerization. Cells were then pelleted and washed in human organoid wash media (Advanced DMEM/F12, 10mM HEPES, 1x GlutaMAX, 100µg/mL Primocin, 0.1% BSA) prior to pelleting again. Cell pellets were passaged at a ratio of 1:2 and replated in Matrigel in new 24-well plates and placed in the incubator for 10 minutes to allow Matrigel to harden. 500µl Human Complete Feeding Media was added on top of Matrigel domes and plates were returned to the incubator for further expansion.

CAFs were extracted from surgical resection specimens after remnant tissue was washed twice with human organoid wash media and strained through a 70µm cell strainer. Cells were plated into 1 well of a 6-well plate and allowed to expand before further cell passaging and expansion. CAFs were expanded by trypsinization and expanded at a ratio of 1:2 in CAF media (RPMI [Fisher 11-875-085] + 10% FBS + 1% Pen/Strep [Gibco 5140122] + 1% L-Glutamine [Gibco 25030081] + 0.1% Amphotericin B [Sigma A2942]). Both cell types were Mycoplasma tested using Invivogen MycoStrips (rep-mys-100) upon establishment and at 6 month intervals thereafter.

#### Coculture setup and cell acquisition

Prior to coculture setup, PDOs and CAFs were lineage verified using short tandem repeat evaluation completed at the Johns Hopkins Genetics Resource Core Facility (Supplemental Table 3). CAFs were grown in monolayer and characterized with flow cytometry using common CAF markers such as VIM, PDPN, PDGFRα, FAP, and αSMA to demonstrate preserved interpatient heterogeneity (Supplemental Figure 6A-C). Epithelial PDOs were established and expanded from primary PDAC tumors as previously described^57^. PDOs were confirmed to be viable, and EpCAM positive (Supplemental Figure 6D-F). Further CAF characterization was performed, in line with prior studies, as cell type markers can display high levels of variability^11,18,59,60^. To more discretely examine CAF heterogeneity and demonstrate specificity of cell type marker expression, we used qPCR to examine gene expressison of 7 CAF and 5 tumor markers in 8 represetative CAF lines and 6 representative PDOs (Supplemental Figure 6G). This allowed us to better define the CAF subtypes prior to inclusion in a three dimensional organotypic co-culture system.

We further optimized a novel PDO-CAF coculture method by taking a reductionist approach and considering the effects of media composition on resulting cell counts (Supplemental Figure 7A-C; Supplemental Table 2) and Matrigel dissociation methods on cell yield (Supplemental Figure 7D) prior to proceeding with further coculture studies. We also compared the effects of different PDO-CAF ratios on CAF expression of αSMA (Supplemental Figure 7E-F); PDOs and CAFs were combined at ratios of 1:1, 1:2, 1:3, 1:5, and 1:10 organoids to CAFs. Cocultured cells were resuspended in different concentrations of Matrigel from 100% to 25% Matrigel (Corning, 356234). To generate lower Matrigel content conditions, Matrigel was supplemented with 5% CAF Media (RPMI + 5% FBS + Pen/Strep, 0.1% Amphotericin B) and plated in triplicate in 24-well tissue culture dishes. Matrigel domes were allowed to harden at 37℃ for 1 hour before adding 500µl CAF media containing 5% FBS to overlie the dome. To extract cells from coculture, supernatant was aspirated, and each dome was digested with either 1mg/mL dispase solution or 2mg/mL collagenase IV for 45min at 37℃. Wash media was added to each well to quench collagenase digest and each well was transferred to a 96-well deep well plate to collect. The plate was centrifuged for 5 minutes at 1500 RPM, supernatant was aspirated, and cells were resuspended in wash media.

Centrifugation and supernatant aspiration were then repeated. Cell pellets were resuspended in TrypLE Express (ThermoFisher Scientific, 12604013) following manufacturer instructions to dissociate organoids into single-cell suspension for use in downstream assays.

### RNA seq

#### Sample preparation and alignment

RNA sequencing sample preparation, library construction, quality control, sequencing, and alignment was done as previously described in detail by Guinn and colleagues^56^. Briefly, cocultured cells were isolated, sorted by FACS and immediately underwent RNA extraction (Qiagen RNeasy Kit, 74004). Quality was assessed by Nanodrop1000 (Thermo Fisher Scientific) and via an external Novogene assessment using an Agilent 2100 Bioanalyzer (Agilent, Santa Clara, CA). mRNA was purified using a magnetic bead approach and cDNA was constructed using random hexamer primers. Libraries were checked with Qubit and pooled for sequencing Illumina platforms to a depth of 40 million reads per sample. The co-cultured organoid sample for JHH 390 did not meet quality metrics upon sequencing, so it was held from analysis. All downstream analysis that include a monoculture to coculture comparision exclude JHH 390.

Alignment was performed utilizing Salmon v1.9.0^61^ on the Joint High-Performance Computing Exchange (JHPCE). “salmon_partial_sa_index__default.tgz” was used as the index for alignment for the HG38 genome, which was premade and available on refgenie^62^.

#### Differential gene expression and pathway analysis

We evaluated sample quality from the distribution of reads as visualized in a boxplot of log counts. We observed no samples with zero median expression, reflective of a low read count, so all samples have good quality. We used principal component analysis (PCA) of the variance stabilization transform (vst) RNA-seq data to evaluate sample clustering. Two samples were identified as not expressing canonical markers of their respective cell types, and being these were of the same patient and timepoint, a renaming was completed to correct for this. Differential expression analysis was completed multiple ways for this study using the DESeq2^63^ package (v1.32.0): coculture organoids compared to monoculture organoids and coculture CAFs compared to monoculture CAFs.

Estimated fold changes were shrunk with apeglm^64^ using lfcShrink to account for the variation in the samples in this dataset. Genes were statistically significant if the absolute log2-fold changes after shrinkage were greater than 0.5 and the FRD-adjusted p-values below 0.05. Gene set statistics were run with fgsea^65^ using MSigDb v7.4.1^66^ pathways annotated in the HALLMARK, KEGG, REACTOME, ONCOGENIC and GO databases. Gene sets were considered to be significantly enriched with FDR-adjusted p values below 0.05. The results were visualized with ggplot2^67^.

### CibersortX

To better understand the cellular heterogeneity in the PDOs and CAFs, raw counts were uploaded to CIBERSORTx^68^ for further deconvolution of cell subtypes. Imputation of cell fractions was completed using a signature matrix generated from reference single-cell RNA-seq data of PDAC tumors which we previously collated and standardized from a range of public domain datasets^23^. This allowed for the estimation of cell fraction based on the bulk gene expression using the default parameters and 1000 permutations for statistical analysis. Results were plotted on a heatmap utilizing ComplexHeatmap^69^ v2.8.0 and comparison boxplots using ggplot2.

## Data and code availability

Raw reads for the RNA-seq data are being submitted to dbGAP and processed read counts are being submitted to GEO. IMC data are available at Zenodo (10.5281/zenodo.14219608). All scripts from analysis will be made available through Github.

## Acknowledgements

This work was funded by the The Lustgarten Foundation (EMJ), NIH/NCI (U01CA253403 to EJF; P01CA247886 to EMJ; S10OD034407 to WJH; K08CA248710 to RAB), the Charles and Margaret Levin Family Foundation, the Dana & Albert R. Broccoli Charitable Foundation, and the MD Anderson GI SPORE. D.J.Z. is supported by a MacMillan Pathway to Independence Award, the MD Anderson GI SPORE Career Enhancement Award, and the Maryland Cancer Moonshot Research Grant to the Johns Hopkins Medical Institutions (FY24). We would also like to thank The Sidney Kimmel Comprehensive Cancer Center Oncology Tissue Services, and the Johns Hopkins Reference Histology Lab and Genetics Resources Core Facility. Finally, we are grateful to the patients who have generously consented to tissue acquisition for this work.

---

**Supplemental Table 1.** IMC markers.

| **<u>Mass</u>** | **<u>Metal</u>** | **<u>Antigen</u>** | **<u>Clone</u>** | **<u>Dilution</u>**<br>**<u>Factor</u>** | **<u>Source</u>** | **<u>Custom</u>** |
| --- | --- | --- | --- | --- | --- | --- |
| 89 | Y | CD45 | D9M8I | 125 | Cell Signaling Technology® | X |
| 96-104 | Ru | Counterstain |  |  | Electron Microscopy Sciences |  |
| 113 | In | Collagen | E8F4L | 250 | Cell Signaling Technology® | X |
| 115 |  | E-Cadherin | 24E10 | 125 | Cell Signaling Technology® | X |
| 141 | Pr | αSMA | 1A4 | 500 | Standard BioTools™ | |
| 142 | Nd | Podoplanin | D2-40 | 125 | Biolegend® | X |
| 143 | Nd | Vimentin | D21H3 | 500 | Standard BioTools™ |  |
| 144 | Nd | S100A4 | D9F9D | 250 | Cell Signaling Technology® | X |
| 145 | Nd | CD45RO | UCHL1 | 250 | Biolegend® | X |
| 146 | Nd | CD16 | EPR167 84 | 100 | Standard BioTools™ |  |
| 147 | Sm | CD163 | EDHu-1 | 125 | Standard BioTools™ |  |
| 148 | Nd | Pan-Keratin | C11 | 125 | Standard BioTools™ |  |
| 149 | Sm | TFF1/pS2 | D2Y1J | 200 | Cell Signaling Technology® | X |
| 150 | Nd | PD-L1 | E1L3N | 125 | Cell Signaling Technology® | X |
| 151 | Eu | PD-1 | D4W2J | 125 | Cell Signaling Technology® | X |
| 152 | Sm | CD31 | 89C2 | 250 | Cell Signaling Technology® | X |
| 153 | Eu | Tox/Tox2 | E6I3Q | 250 | Cell Signaling Technology® | X |
| 154 | Sm | CD57 | HNK-1 | 250 | Cell Signaling Technology® | X |
| 155 | Gd | Foxp3 | PCH101 | 75 | Standard BioTools™ |  |
| 156 | Gd | CD4 | EPR685 5 | 125 | Standard BioTools™ |  |
| 158 | Gd | N-Cadherin | D4R1H | 125 | Cell Signaling Technology® | X |
| 159 | Tb | CD68 | KP1 | 100 | Standard BioTools™ |  |
| 160 | Gd | IL-6 | D3K2N | 125 | Cell Signaling Technology® | X |
| 161 | Dy | CD20 | H1 | 125 | Standard BioTools™ |  |
| 162 | Dy | CD8α | C8/144 B | 250 | Standard BioTools™ |  |
| 163 | Dy | PDGFRα | D1E1E | 250 | Cell Signaling Technology® | X |
| 164 | Dy | IL-8 | E5F5Q | 250 | Cell Signaling Technology® | X |
| 165 | Ho | GATA6 | Polyclo<br>nal | 250 | R&D Systems™ | X |
| 166 | Er | CD45RA | HI100 | 250 | Standard BioTools™ |  |
| 167 | Er | Granzyme B | D6E9W | 125 | Cell Signaling Technology® | X |
| 168 | Er | Ki-67 | B56 | 250 | Standard BioTools™ |  |
| 169 | Tm | CXCL12 | D8G6H | 125 | Cell Signaling Technology® | X |
| 170 | Er | CD3ε | Polyclo<br>nal, C-termina<br>l | 125 | Standard BioTools™ |  |
| 171 | Yb | S100A2 | Polyclo<br>nal | 125 | Novus Biologicals™ | X |
| 172 | Yb | FAPα | EPR200<br>21 | 250 | Abcam | X |
| 173 | Yb | CD137 (4-1BB) | D2Z4Y | 125 | Cell Signaling Technology® | X |
| 174 | Yb | HLA-DR | LN3 | 250 | Standard BioTools™ |  |
| 175 | Lu | CD105 | 3A9 | 125 | Cell Signaling Technology® | X |
| 176 | Yb | KRT17 | D12E5 | 250 | Cell Signaling Technology® | X |
| 191 | Ir | DNA 1 |  |  | Standard BioTools™ |  |
| 193 | Ir | DNA 2 |  |  | Standard BioTools™ |  |
| 195 | Pt | Plasma Membrane 2 | 1A36 | 250 | Standard BioTools™ |  |
| 196 | Pt | Plasma Membrane 3 | 1A37 | 250 | Standard BioTools™ |  |
| 198 | Pt | Plasma Membrane 4 | 1A38 | 250 | Standard BioTools™ |  |

**Supplemental Table 2.** HCPLT media.

| **<u>Volume</u>** | **<u>Stock Concentration</u>** | **<u>Reagent</u>** | **<u>Final Concentration</u>** |
| --- | --- | --- | --- |
| 36.415 mL |  | Human Wash Media |  |
| 50 mL | 2X | Wnt2a-conditioned media | 1X |
| 10 mL | 10X | R-spondin1-conditioned media | 1X |
| 2 mL | 50X | B27 supplement | 1X |
| 1 mL | 1 M | Nicotinamide | 10mM |
| 250 uL | 500 mM | N-acetylcysteine | 1.25mM |
| 200 uL | 50 mg/mL | Primocin | 100ug/mL |
| 100 uL | 100 ug/mL | mNoggin | 100ng/mL |
| 10 uL | 500 ug/mL | hEGF | 50ng/mL |
| 10 uL | 1000 ug/mL | hFGF | 100 ng/mL |
| 10 uL | 100 uM | hGastrin I | 10 nM |
| 2 uL | 25 mM | A 83-01 | 500 nM |
| 100 uL | 10.5 mM | Y-27632* | 10.5 uM |

Y-27632 (Rho Kinase Inhibitor) is necessary to maintain organoids after first generation, when organoids are thawed, or when organoids are enzymatically dissociated.

**Supplemental Table 3.** Lineage Verification.

| **<u>PDO Line</u>** | **<u>%Match</u>** | **<u>Alleles shared</u>** | **<u>Alleles in line 1 (CAF)</u>** | **<u>Alleles in line 1 (org)</u>** |
| --- | --- | --- | --- | --- |
| JHH317 | 97.14% | 17 | 17 | 18 |
| JHH348 | 61.11% | 11 | 17 | 19 |
| JHH352 | 96.97% | 16 | 17 | 16 |
| JHH357 | 100% | 18 | 18 | 18 |
| JHH361 | 84.85% | 14 | 19 | 14 |
| JHH368 | 97.14% | 17 | 18 | 17 |
| JHH369 | 100% | 19 | 19 | 19 |
| JHH372 | 94.44% | 17 | 19 | 17 |
| JHH380 | 100% | 16 | 16 | 16 |
| JHH383 | 100% | 16 | 16 | 16 |
| JHH387 | 100% | 19 | 19 | 19 |
| JHH390 | 100% | 15 | 15 | 15 |

**Supplemental Table 4.**
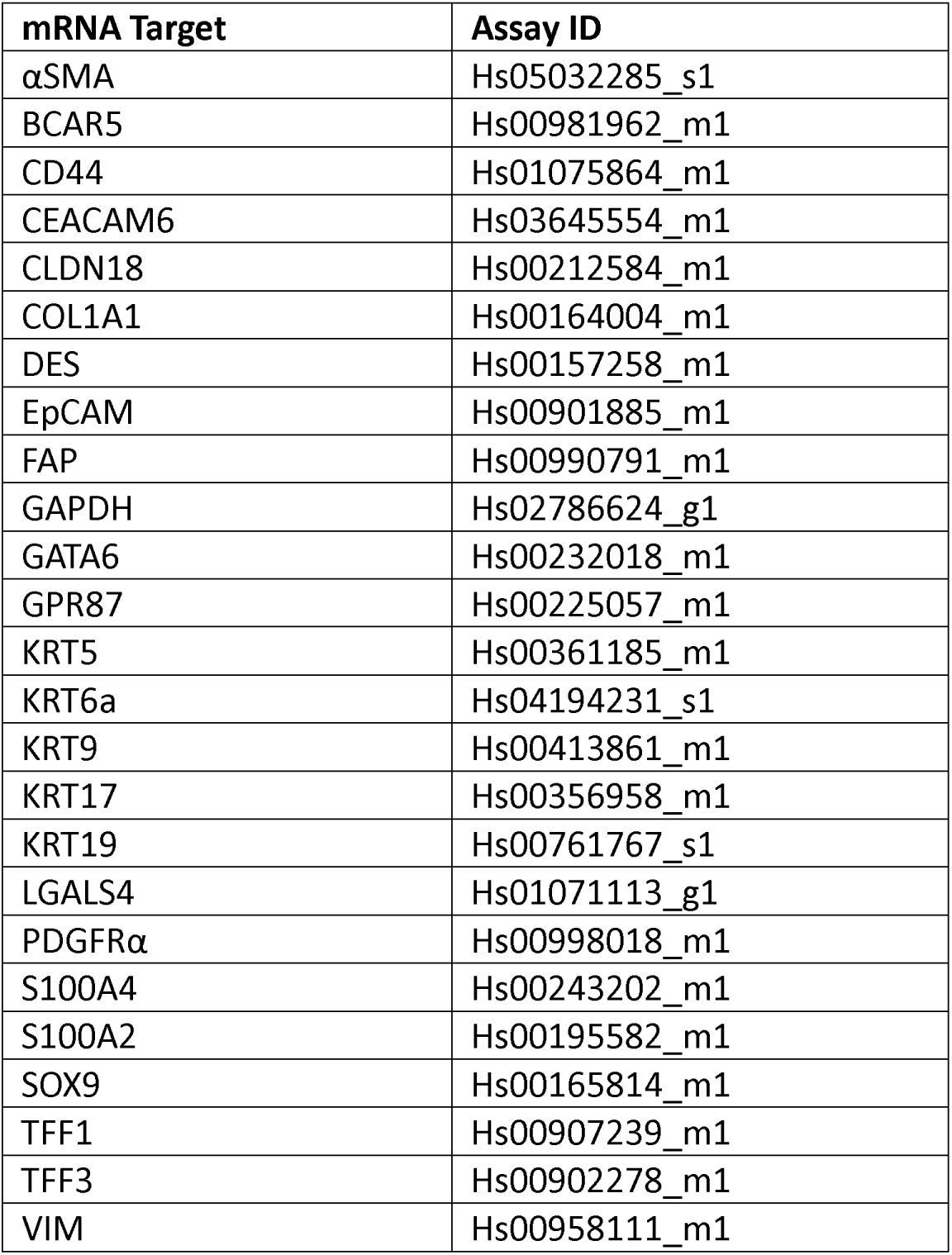
mRNA Target Assay ID.

**Supplemental Figure 1.**
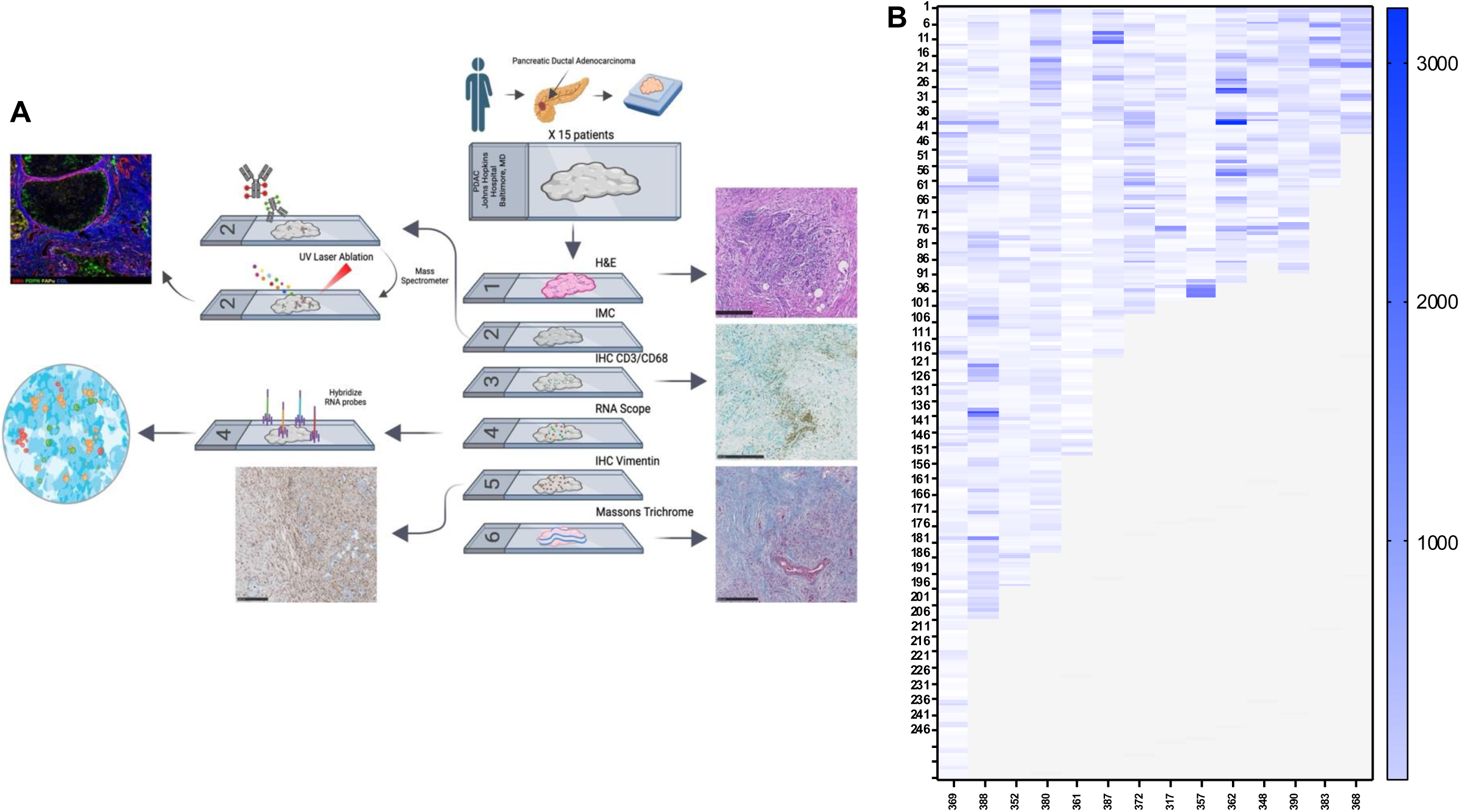

**Supplemental Figure 2.**
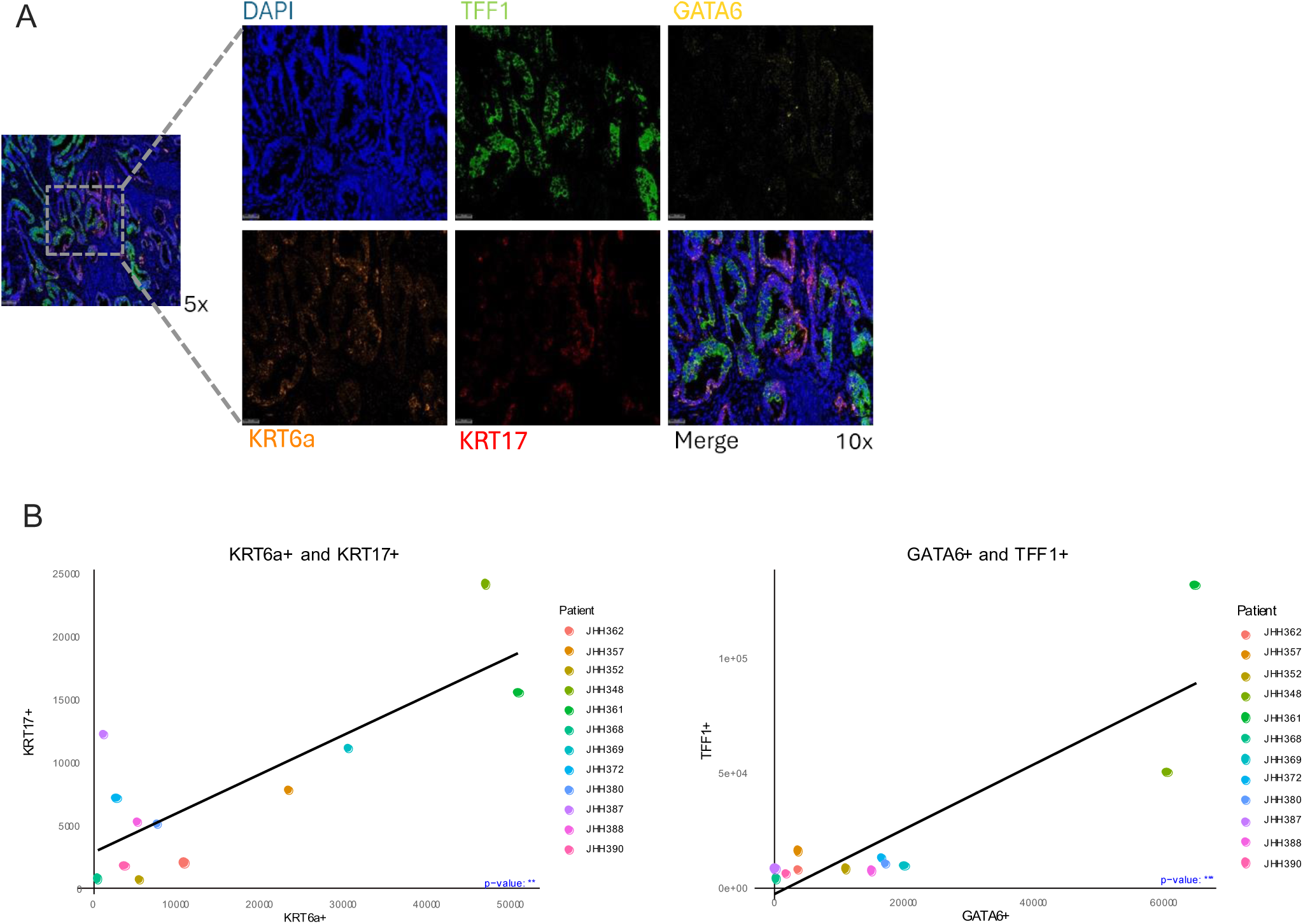

**Supplemental Figure 3.**
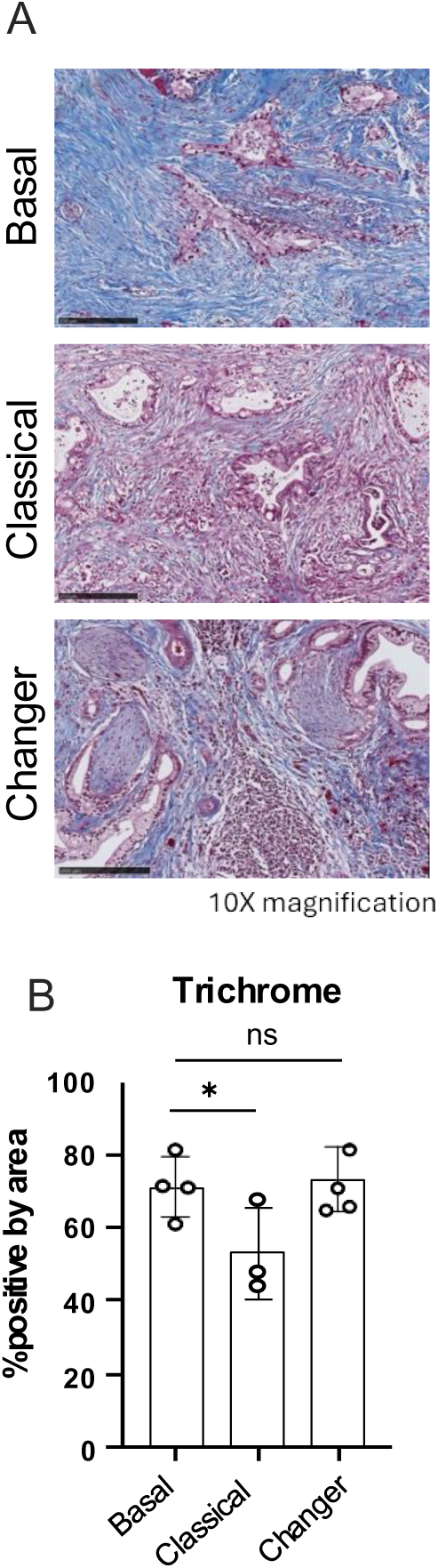

**Supplemental Figure 4.**
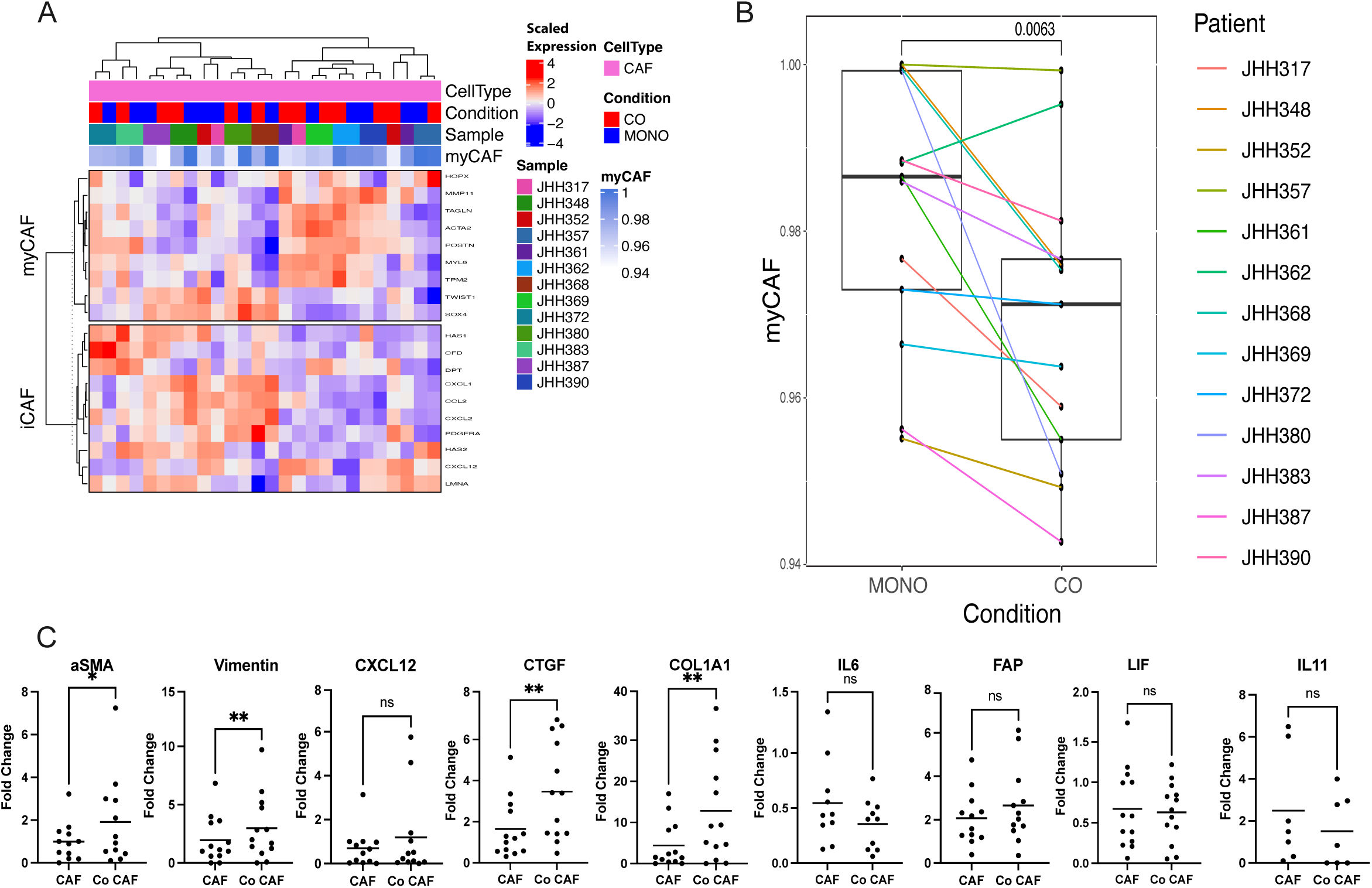

**Supplemental Figure 5.**
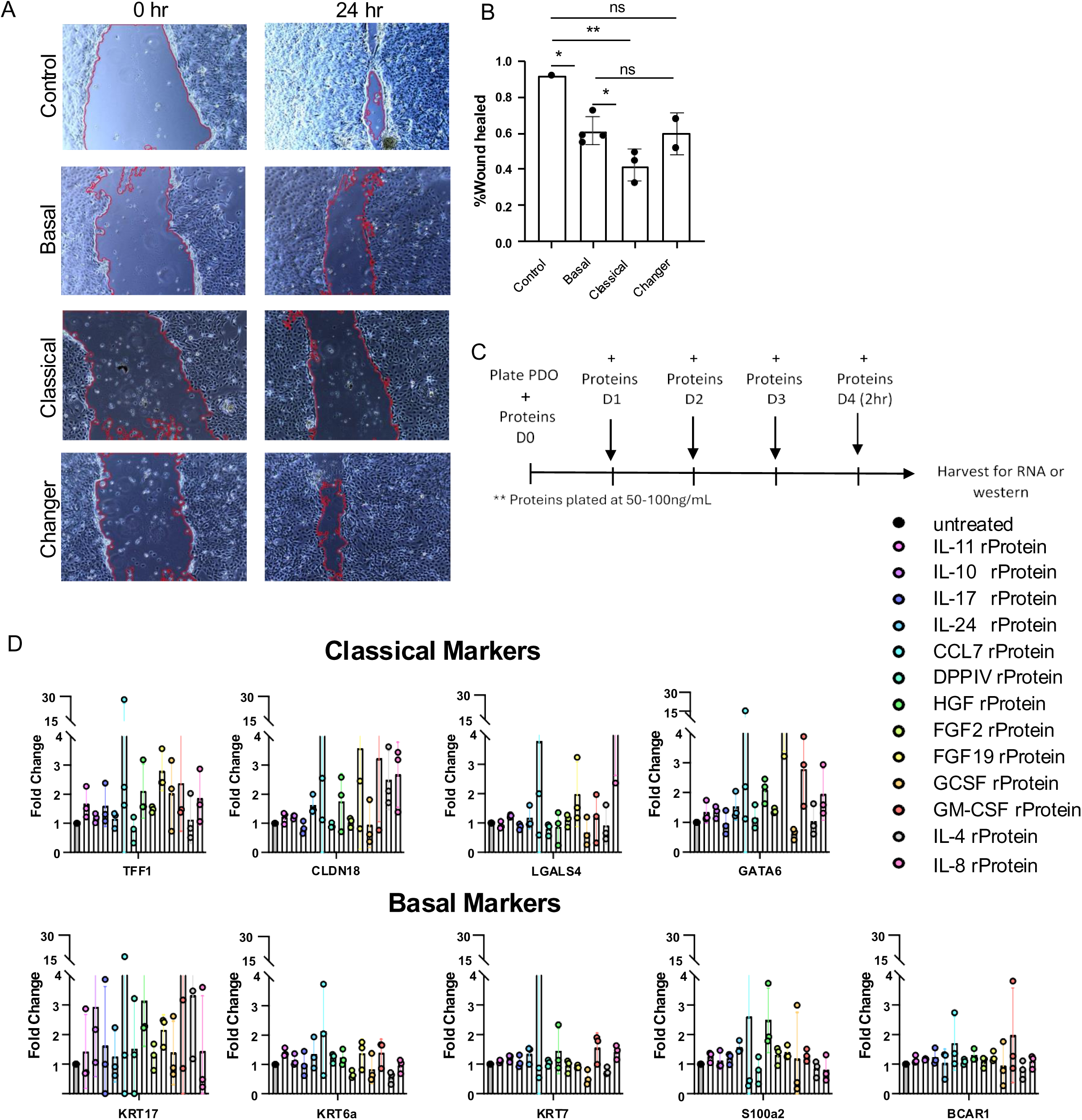

**Supplemental Figure 6.**
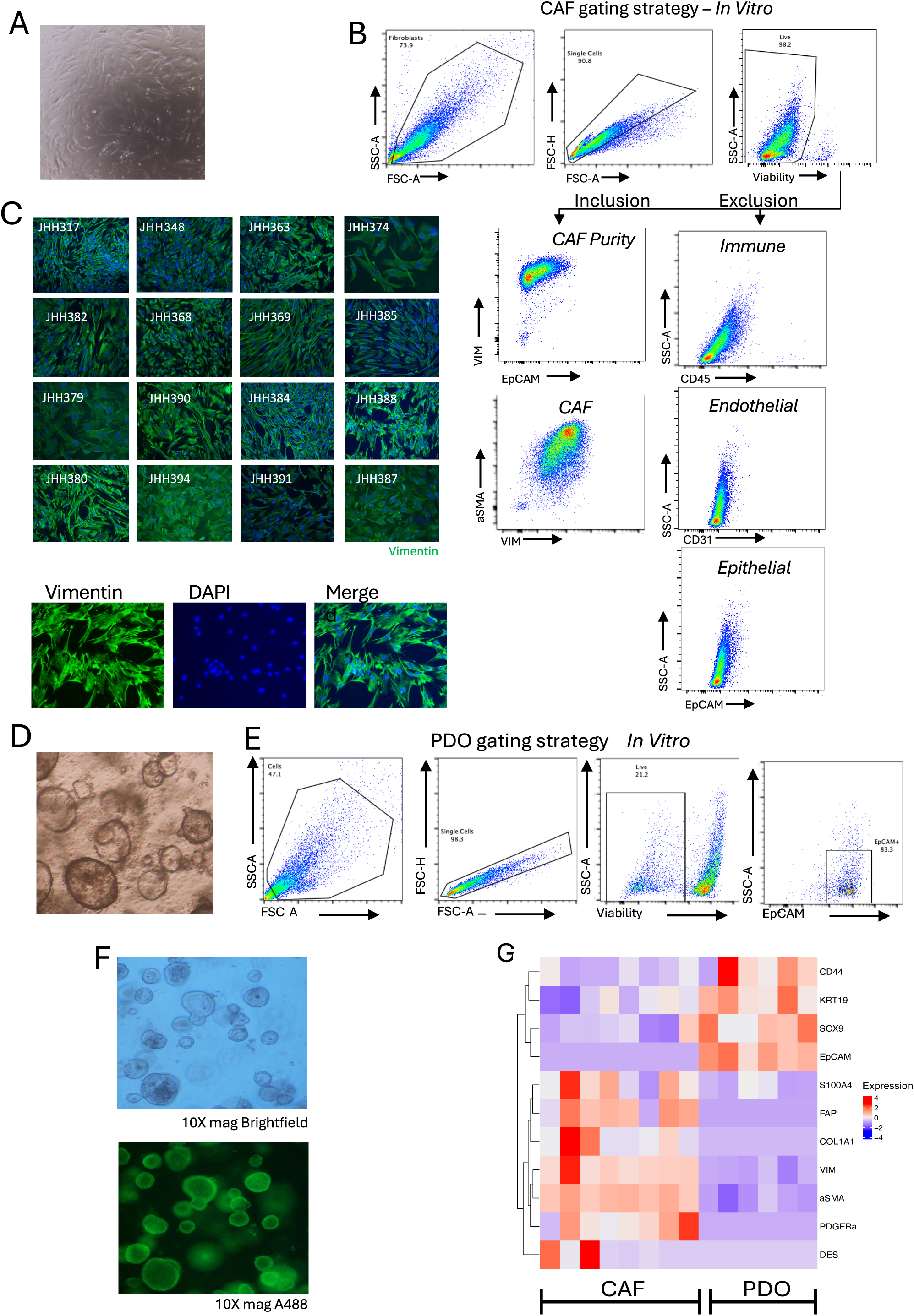

**Supplemental Figure 7.**
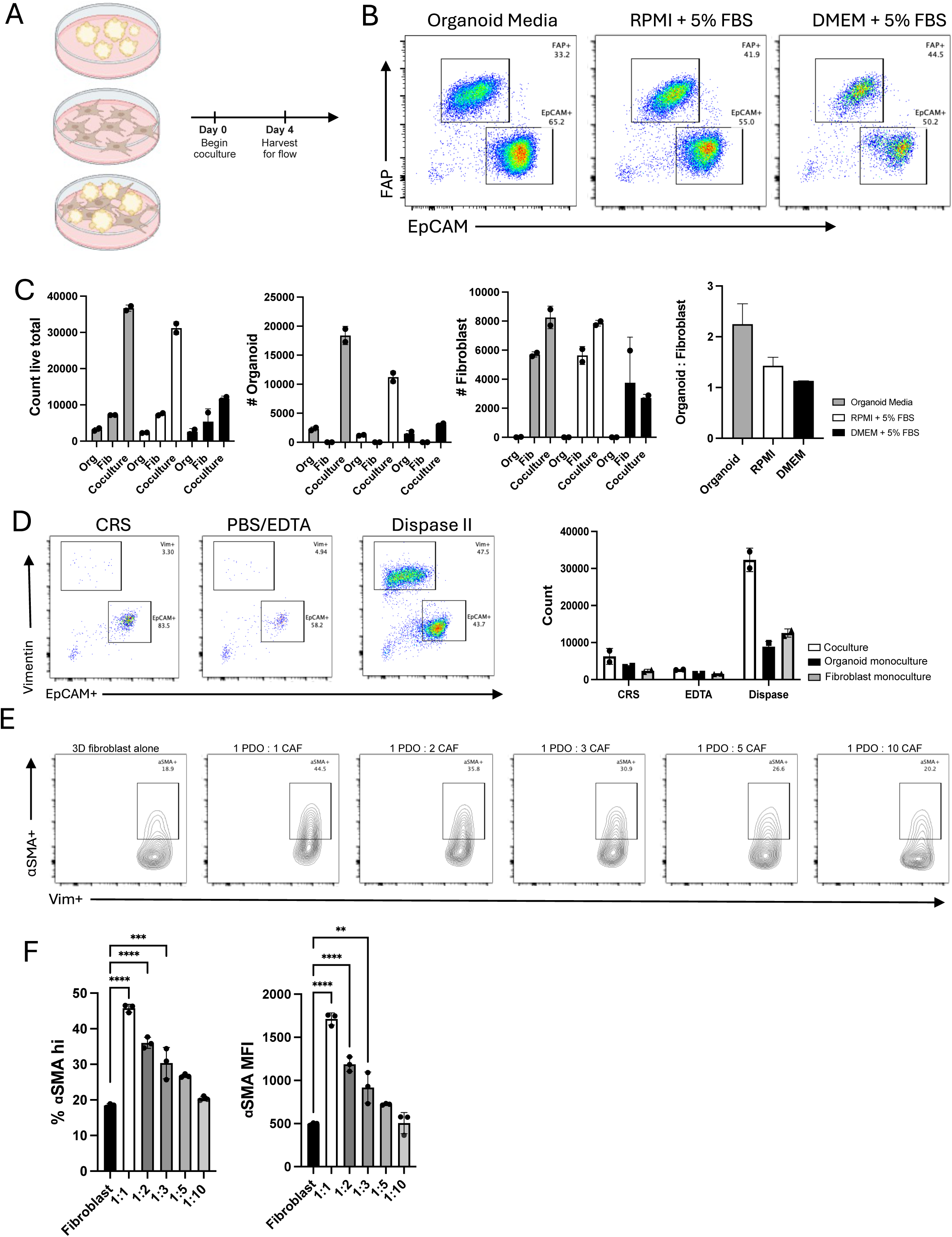

