## Supplemental Figure Legends for "Cancer associated fibroblasts drive epithelial to mesenchymal transition and classical to basal change in pancreatic ductal adenocarcinoma cells with loss of IL-8 expression"

**Supplemental Figure 1.** (A) Serial sections were cut from blocks chosen by an expert pathologist containing 4 regions of interest for analysis using imaging IMC, IHC, and RNAscope. (B) Ranking system of CD3+ staining per patient. Each patient slide was analyzed using a grid system 1µm^2^ squares in order to calculate how many CD3+ cells were present per 1µm^2^ square

**Supplemental Figure 2.** (A) (Far left) RNA Scope representative image of patient tissue at 5X magnification and a grid analysis system applied to analyze the tissue regions. (Top left) DAPI (blue), (Top middle) TFF1 (Green), (Top right) GATA6 (Yellow), (Bottom left) KRT6a (orange), (Bottom middle) KRT17 (red), and (Bottom right) merge (all channels). (B) (Left) Scatterplot correlation between basal markers (top) KRT17 on y axis, and KRT6a on x axis for each patient (Right) Scatterplot correlation between classical markers (top) TFF1 on y axis, and GATA6 on x axis for each patient.

**Supplemental Figure 3.** (A) Trichrome image representation of patient tissue for one of the patients in this study at 10X magnification (scale bar = 250µm). Patients chosen were from those classified as either basal, classical, or "changer" by RNA sequencing designation. (B) Quantification of tumor regions in (A). Statistics done by students T test, * = p <0.05.

**Supplemental Figure 4.** (A) Heatmap displaying myCAF and iCAF gene expression in CAFs between coculture and monoculture samples. (B) myCAF gene expression on a per patient basis. Comparisons of conditions are statistically supported using the two-tailed students t-test with equal variance p=0.0063. (C) qPCR expression of monoculture CAF and coculture CAF for 9 different genes. Comparisons of conditions are statistically supported using the two-tailed students t-test with equal variance. Significance is measured as: ****, p<0.0001; ***, p<0.001; **, p<0.01; *, p<0.05; ns, not significant.

**Supplemental Figure 5.** (A) Representative brightfield images from a scratch wound healing assay for Panc10.05 cells treated as control (Panc10.05 cells with untreated CAF media), or treated with CAF conditioned media from patient lines that were classical, basal, or “changer”. Wound edge outlined in red. (B) Quantification of percentage of time 0 healed at 24hr from (A). Three technical replicates per fibroblast conditioned media group. (C) Experimental setup timeline of recombinant protein screen for 14 proteins that were queried. (D) Classical (top) and basal (bottom) exploration for each of the 14 proteins screened relative to untreated (black, far left bar). Three or four when available, technical replicates per target. Significance is measured as: ****, p<0.0001; ***, p<0.001; **, p<0.01; *, p<0.05; ns, not significant.

**Supplemental Figure 6.** (A) Representative image of CAFs grown in 2D on tissue culture treated flasks for expansion and downstream experiments. (B) Flow cytometry gating scheme of CAFs harvested from these flasks to ensure there were no contaminating cells, negative gate including CD45 for immune cells, CD31 for endothelial cells, and EpCAM for epithelial cells. Positive CAF selection by a combination of VIM, FAP, and αSMA. (C) Top, immunofluorescence staining for vimentin in CAFs. Variation of VIM+ cells was detected per patient, and patient heterogeneity was preserved. Bottom, Individual channels of the immunofluorescence (C) showing vimentin in GFP (far left), DAPI in blue (middle), and merge (far right). (D) Representative image of PDOs grown in 3D in Matrigel and expanded and maintained over time. (E) Flow plots showing sustained PDO gating strategy and EpCAM+ expression on the organoids. (F) (top) Brightfield imaging of PDO at 10x magnification, (bottom) EpCAM+ expression of PDO (a488 staining at 10x magnification). (G) Heatmap showing expression of PDO and CAF markers by qPCR of 11 targets investigated.

**Supplemental Figure 7.** To successfully create a novel multicompartmental coculture, one major consideration was that PDOs are routinely cultured in mitogen rich media and CAFs are cultured in their own CAF media. The additives in the PDO media include TGFβ, noggin, and other inhibitors of CAF proliferation (Supplemental Table 2). (A) Experimental schema of coculture to further vet the media composition and readout. (B) Organoid media, RPMI + 5% FBS, and DMEM + 5% FBS were tested in coculture to determine optimal media composition. Both PDOs and CAFs could be cultured in low serum RPMI media for the duration of coculture. (C) Quantification of media test in (B). (D) Methods tested to optimize extraction method to select for highest number of CAFs and PDOs to be extracted from Matrigel. Conditions tested include Cell Recovery Solution (CRS) (Left), PBS + 0.5mM EDTA (middle), and 1mg/mL Dispase II (right). Quantification shown by total cell count recovered (far right). Dispase II liberated the greatest number of CAFs when compared to Cell Recovery Solution or a combination of PBS and EDTA. (E) Evaluation of cell ratios (and the resulting differences in cell proximity) in determining cell phenotype by coculturing PDOs and CAFs at increasing CAF ratios. Increased CAF presence induced a myCAF-like phenotype, as determined by upregulation of αSMA. Flow plots demonstrate Vim and αSMA expression in cocultured and monocultured CAFs at decreasing ratios of PDOs to CAFs. (F) Quantification of (E) by overall percentage change in αSMA expression (left), or by αSMA MFI change (right). There was a significant increase in CAF αSMA expression at a ratio of 1 PDO: 3 CAFs as compared to higher CAF ratios, and therefore moved forward with this ratio for the remaining experiments Comparisons of ratios are statistically supported using a two-way ANOVA in PRISM (V9.2.0 [283]). Significance is measured as: ****, p<0.0001; ***, p<0.001; **, p<0.01; *, p<0.05; ns, not significant.
