## Supplemental Methods for "Cancer associated fibroblasts drive epithelial to mesenchymal transition and classical to basal change in pancreatic ductal adenocarcinoma cells with loss of IL-8 expression"

**Quantitative PCR (qPCR)**

To evaluate gene expression, total RNA was extracted from PDO and CAF lines using the RNeasy Mini Kit (Qiagen, 74104) according to manufacturer specifications. cDNA synthesis was performed using TaqMan Reverse Transcription Reagents (Invitrogen, N8080234), following manufacturer’s instructions. Real-time quantitative PCR was completed using the ThermoFisher Taqman Gene Expression Assays according to manufacturer’s protocol in the QuantStudio 6 Flex System (Applied Biosystems). mRNA targets are listed in Supplemental Table 4. Relative gene expression was quantified using the 2^−ΔΔCt^ method as previously described ^1^, and GAPDH was used as the endogenous control. Data were analyzed using Applied Biosystems QuantStudio^TM^ Real Time PCR System Software (V1.7.1).

**Flow Staining and Cell Sorting**

Organoids were extracted using Cell Recovery Solution (Corning, 354253) and incubated on ice at 4°C for 45 minutes for Matrigel depolymerization. Cells were then pelleted and washed in human organoid wash media. Organoids were dissociated to single cells using TrypLE Express and washed in MACS buffer (PBS + 5 mM EDTA + 1% FBS). Cells were then resuspended in PBS + Zombie NIR (BioLegend 423106; dilution 1:1000) + Human TruStain FcX (BioLegend, 422302; dilution 1:100) for 10 minutes at room temperature in the dark. Cells were then quenched with MACS buffer and spun down. Cells were resuspended in surface stain for 20 minutes on ice at 4°C. Cells were washed twice in MACS buffer. Flow cytometry analyses were performed on Beckman Coulter Cytoflex.

For coculture cell sorting, 1mL of 1mg/mL Dispase II in wash media (Thermofisher, 17105041) was added to each coculture and monoculture dome to depolymerize Matrigel for 1 hour at 37°C. Digest was quenched with 1mL of wash media and cells were spun down. Cells were resuspended in PBS + Zombie NIR (BioLegend, 423106; dilution 1:1000) + Human TruStain FcX (BioLegend, 422302; dilution 1:100) for 10 minutes at room temperature in the dark. Cells were quenched with MACS buffer, centrifuged, and then resuspended in surface stain for 20 minutes on ice at 4°C in the dark; PE FAP (R&D, FAB3715P; dilution 1:75), APC EpCAM (BioLegend, 324208; dilution 1:200), Cells were washed twice in MACS buffer. Flow cytometry was performed on the Beckman Coulter Cytoflex and analyzed by FlowJo version 10.8.0.

**Lentiviral transduction**

Lentiviral production was done as described in the protocol developed by the TRC library (Broad Institute). Briefly, HEK293T cells were co-transfected with a gene expression plasmid and second generation lentiviral packaging plasmids (psPAX2, Addgene 12260; VSV-G Addgene 8454; pLV[Expression]-mCherry:T2A:Hygro-EF1A>Luc2; Addgene, 174665). Media was changed the next day and the supernatant-containing virus was harvested at 48 hours, filtered through a 0.45 μm low protein binding PES filter. Lentivirus was concentrated with Concentration Solution (GeneCopoeia, LPR-LCS-01). Optimal transduction units were 4.93X10^7^. Target cells were plated with 250µl of concentrated lentivirus containing media supplemented with 1µl of 1mg/mL hexadimethrine bromide (Sigma-Aldrich H9268) and spin transfected at 1700 RPM for 60 minutes at room temperature. Plates were incubated for 1-6 hours. Cells were resuspended in wash media and spun down and plated into Matrigel domes. 500µl of HCPLT media + Rho Kinase inhibitor (dilution 1:1000) were added, and plates were incubated for 48 hours. Cells were selected by either hygromycin at 200µg/mL or by flow sorting on a FACSAria Fusion (BD) for the selection marker (mCherry).

**Slide Stains**

CD3/CD68 dual immunohistochemistry (IHC) staining

Immunostaining was performed at the Sidney Kimmel Comprehensive Cancer Center Oncology Tissue Services Core. Dual chromogenic immune-labeling for CD3 and CD68 was performed on formalin‐fixed, paraffin embedded sections on a Ventana Discovery Ultra autostainer (Roche Diagnostics). Briefly, following dewaxing and rehydration, epitope retrieval was performed using Ventana Ultra CC1 buffer (6414575001, Roche Diagnostics) at 96^o^C for 64 minutes. Primary antibody, anti‐CD3 (Abcam, ab16669; dilution 1:200) was applied at 36^o^C for 60 minutes and detected using an anti-rabbit HQ detection system (Roche Diagnostics, 7017936001 and 7017812001) followed by Chromomap DAB IHC detection kit (Roche Diagnostics, 5266645001). Primary antibodies were stripped at 95^o^C for 12 minutes and residual HRP was neutralized (Roche Diagnostics, 7017944001). Next, primary antibody CD68 (Dako, M0814; dilution 1:5000) was applied at 36^o^C for 60 minutes and detected using an anti-mouse HQ detection system (Roche Diagnostics, 7017936001 and 7017782001) followed by Discovery Teal Detection kit (Roche Diagnostics, 8254338001), counterstaining with hematoxylin, bluing, dehydration and mounting. Slides were imaged using a Hamamatsu NanoZoomer digital slide scanner at 20x magnification using the associated NDP.toolkit slide processing software. Slides were quantified by tiling the tumor area in a grid with individual regions set to a 1µm^2^ to account for the size of imaging mass cytometry ROIs and analyzed for total cell count utilizing HALO Multiplex IHC v3.4.9 software (Indica Laboratories)

Single target IHC staining

Paraffin-embedded patient tumors were sectioned at 5µm thickness and tissue was rehydrated through a series of xylene and alcohol washes, which were followed with a rinse in water and PBS wash. Slides were put in an antigen retrieval buffer (3300 Vector Labs, 3300) and steamed for 20 minutes. Slides were then blocked in a peroxide blocking buffer (Abcam, ab64218) for 15 minutes, followed by protein block (Abcam, ab64226) for 5 minutes. They were then incubated with anti-Vimentin primary antibody (Cell Signaling Technology, 5741S; dilution 1:200), which was prepared in antibody diluent (Dako, S0809). Slides were placed in a humidified chamber at 4°C overnight. The next day slides were washed with PBS and incubated in biotinylated anti-rabbit (Abcam, ab64256), followed by streptavidin–HRP solution (Abcam, ab64269) at room temperature for 20 minutes. Samples were then washed with PBS and incubated with AEC (3-amino-9- ethylcarbazole) chromogen (Vector Laboratories, SK-4200). Slides were then washed with water and incubated in Mayer’s hematoxylin (Sigma, MHS1) for 1 minute, rinsed with water, and mounted in VECTASHIELD PLUS (Vector Laboratories, H-1900). Slides were imaged using a Hamamatsu NanoZoomer digital slide scanner at 20x magnification using the associated NDP.toolkit slide processing software.

Slides were quantified by tiling the tumor area in a grid with individual regions set to a 1µm^2^ to account for the size of imaging mass cytometry ROIs and analyzed for total cell count utilizing HALO Multiplex IHC v3.4.9 software (Indica Laboratories).

Trichrome staining

Staining was performed by the Johns Hopkins Reference Histology Core. Analysis was performed using the colour_deconvolution2 plugin for ImageJ. Images were threshold adjusted on the blue image to adjust the threshold covering all regions of collagen that were seen and analyzed through ImageJ. Defined regions of tumor rich tissue were quantified and analyzed in Prism.

H&E staining

H&E staining was performed by the Johns Hopkins Reference Histology Core.

**Immunofluorescence**

CAF cells were grown to 80% confluence in 96-well flat bottom tissue treated plates. When desired confluence was reached, media was aspirated and cells were washed twice with PBS. PBS was aspirated and cells were fixed using 10% formalin for 10 minutes. Cells were washed twice more with PBS after fixation was complete. Surface markers were blocked using 5% BSA in PBS for 1 hour at room temperature. Cells were then permabilized with 0.01% Tween for 10 minutes. Primary antibody (VIM, Biolegend, 677809) was added and cells were incubated at 4°C overnight. After incubation, cells were washed twice with PBS and counterstained with DAPI for 30 seconds. DAPI was pipetted off and a small amount of PBS was added to keep the cells wet. Cells were imaged using an inverted microscope with a filter for GFP.

**Wound healing assay**

Pancreatic cancer cells (PANC10.05, ATCC, CRL-2547) were plated at a density of 5x10^4^ cells/well in a 24-well plate and cultured in the indicated media for 48 hours. A 200µL pipette tip was used to introduce a wound into the cell monolayer. Phase contrast images were taken after media was replaced at the time of the wound creation and at the indicated time points. Images were analyzed with Fiji imageJ analysis software to define the area of the wound at time = 0 and again at the end of the experiment. The fraction of wound closure was calculated by dividing the final wound area by the initial wound area and subtracting the resulting value from 1.

**Proteome profiler**

Conditioned media was collected after a 4-day incubation with CAF monoculture, PDO monoculture, or matched cocultures of cell lines that shift from classical to basal, start classical and stay classical, or start basal and remain basal. Media was filtered through a 0.45µm filter to remove cell debris, and frozen at -80°C. Frozen conditioned media was then thawed and used for cytokine dot blots (R&D Systems, ARY022B) according to the manufacturer's instructions. Dot blots were analyzed using Image J with the microarray profile plugin. Normalized Z scores were tabulated and analyzed with Prism V9.5.0.

**In situ hybridization RNAscope**

The FFPE-embedded pancreas tumor tissue sections (5μm thick) were cut onto SuperFrost Plus slides. Staining was completed with an RNAscope Multiplex Fluorescent Reagent kit v2 assay (Advanced Cell Diagnostics, Bio-Techne, 323135). Briefly, target RNAs were hybridized to ssDNA “z-probes” complementary to RNA of interest. Oligos were bound to the tail region of the z-probe, which were then bound to amplifiers labeled with horseradish peroxidase (HRP) and fluorophores. Probes generated by Advanced Cell Diagnostics (520721, c1 – KRT6A; 463661-c2 – KRT17; 603131 – c3, GATA6; 417281- c4, TFF1) were uniquely amplified with Opal dyes (Akoya Biosciences, FP1487001KT, FP1488001KT, FP1497001KT, FP14695001KT) and then counterstained with DAPI (1μg/ml). Slides were mounted using Pro-Long Gold mounting medium (Invitrogen, p10144). Slides were imaged on a Vectra Polaris scanning microscope. Quantification was performed using inForm imaging software (v2.5.1) to demultiplex the sample images and create individual tiles. Tiles were reconfigured and analyzed in HALO using the Highplex FL module software (v4.2.14). The following channels were used: DAPI (excitation, 375 nm; emission, 435–480 nm); Opal 520 (excitation, 488 nm; emission, 500–550 nm); Opal 570 (excitation, 561 nm; emission, 570–630 nm); Opal 620 (excitation, 588 nm; emission, 650–760 nm); Opal 690 (excitation, 676nm, emission, 694nm).
